# Single-cell profiling identifies ACE^+^ granuloma macrophages as a non-permissive niche for intracellular bacteria during persistent *Salmonella* infection

**DOI:** 10.1101/2022.07.21.501041

**Authors:** Trung H. M. Pham, Yuan Xue, Susan M. Brewer, KE Bernstein, Stephen R. Quake, Denise M. Monack

**Author notes:** Contributed equally (Trung H. M. Pham, Yuan Xue). Correspondence and co-lead contacts: Denise M. Monack,; Trung H. M. Pham,; Stephen R. Quake.

## Abstract

Macrophages mediate key antimicrobial responses against intracellular bacterial pathogens, such as *Salmonella enterica*. Yet, they can also act as a permissive niche for these pathogens to persist in infected tissues within granulomas, which are immunological structures comprised of macrophages and other immune cells. We apply single-cell transcriptomics to investigate macrophage functional diversity during persistent *Salmonella enterica* serovar Typhimurium (*S*Tm) infection in mice. We identify determinants of macrophage heterogeneity in infected spleens and describe populations of distinct phenotypes, functional programming, and spatial localization. Using a *S*Tm mutant with impaired ability to polarize macrophage phenotypes, we find that angiotensin converting enzyme (ACE) defines a granuloma macrophage population that is non-permissive for intracellular bacteria and their abundance anticorrelates with tissue bacterial burden. Disruption of pathogen control by neutralizing TNF preferentially depletes ACE^+^ macrophages in infected tissues. Thus ACE^*+*^ macrophages have differential capacity to serve as cellular niche for intracellular bacteria to establish persistent infection.

**Teaser:** This study shows that ACE^*+*^ granuloma macrophages have restricted capacity to act as a cellular niche that enables intracellular bacterial persistence.

## Introduction

Intracellular bacteria such as *Salmonella enterica, Brucella melitensis*, and *Mycobacterium tuberculosis* infect hundreds of millions of people and cause millions of deaths annually(1, 2). These pathogens can establish persistent infection and survive in host tissues at low levels for months to years(3–6). Macrophages mediate key antibacterial immune responses, such as phagocytizing and killing bacteria, producing proinflammatory cytokines, and modulating adaptive immunity(7–9). Yet macrophages also act as a cellular niche for intracellular bacteria to persist within the granulomas, which are tissue microstructures comprised of macrophages and other cell types(10–15). Persistent intracellular bacterial infections pose great clinical challenges due to transmission from asymptomatic carriers, ineffective strategies for monitoring disease progression, and prolonged antimicrobial therapies that increase the risk for developing antimicrobial resistance(16). Modulating the differential functions of macrophages may present a viable therapeutic strategy to limit intracellular bacterial infections. However, our understanding of macrophage heterogeneity and its functional diversity in infected tissues remains largely incomplete.

Although invasive biopsy is not routinely performed, human histopathological studies suggest that macrophage-rich granulomas are a common feature of intracellular bacteria-infected tissues, such as spleens and livers(17–19). Animal infection models have been essential for understanding how diverse tissue macrophage functions and granuloma formation contribute to persistence of bacilli. For example, persistent infection in systemic tissues such as the spleen has been investigated by infecting 129×1/SvJ mice with fully virulent *S. enterica* serovar Typhimurium (*S*Tm)(4, 13, 14, 20, 21). In chronically infected mice, *S*Tm bacilli persist for months at low abundance within splenic granulomas comprised of heterogeneous macrophages. Neutrophils, lymphocytes, as well as other immune and non-immune cells variably contribute to a spectrum of granuloma cellular composition, even among granulomas in the same infected tissue(10, 12, 15). Innate cellular antibacterial functions and T-cell immune responses have been shown to be important for containing pathogens within intracellular bacterial granulomas and controlling infection(12). Recent studies suggest the balance between proinflammatory, M1-like and anti-inflammatory, M2-like macrophage activities within granulomas and intracellular-bacteria infected tissues influences bacterial persistence and eradication(14, 22). How intracellular bacterial pathogens survive within granulomas despite macrophage recognition and antibacterial mechanisms, as well as robust innate and adaptive immune responses remains a central question.

Approaches to defining diverse macrophage phenotypes and functions in infected tissues often depend on biased cellular features derived from a dichotomous paradigm of the classically activated (M1) and alternatively activated (M2) model(23–28). While overly simplified, M1-like macrophages are thought to be pro-inflammatory and antibacterial, whereas M2-like macrophages are thought to be more anti-inflammatory, pathogen permissive, and crucial for tissue repair. Increasing evidence suggests the dichotomous M1 and M2 paradigm is insufficient to fully account for the heterogeneous phenotypes of macrophages *in vivo*. A multitude of factors, including ontogenetic programming, microenvironmental signals, and bacterial effector manipulation shape tissue macrophage heterogeneity and functional diversity(8, 26, 27, 29, 30). Macrophages localizing in different regions of granulomas within intracellular bacteria-infected tissues exhibit distinct phenotypes and may have differential functions such as antibacterial activities and T-cell modulation(13, 14, 31, 32). We and others previously showed that *S*Tm utilizes its *Salmonella* Pathogenicity Island 2 (SPI2) Type 3 Secretion System (T3SS) effector SteE to promote bacterial persistence within granulomas by skewing macrophage phenotype toward a permissive, M2-like state(14, 26, 28). Remarkably, during the persistent infection stage, many granuloma macrophages harboring intracellular *S*Tm also express high levels of the inducible nitric oxide synthase, iNOS, which has been utilized as a canonical marker of M1-like macrophages(13, 14, 33). Tissue macrophages have also been found to have different developmental origins that may influence their capacity to restrict intracellular bacteria. For example, in the lungs of *M. tuberculosis* (*Mtb*)-infected mice, monocyte-derived interstitial macrophages and alveolar macrophages, which originate from embryonic precursor cells, exhibit differential antibacterial capacities(34–36). Despite their ability to restrict intracellular bacilli, alveolar macrophages were shown to exhibit M2-like characteristics and thereby served as a more favorable replicative niche for *Mtb*(27, 36). In a recent single-cell RNA-sequencing (sc-RNAseq) study of acute *S*Tm infection in mice for 48 hours after inoculation, when granulomas are yet to form and bacterial levels are uncontrolled, a non-classical monocyte-derived macrophage population was found to harbor more intracellular bacilli than other mononuclear phagocytes(37).

Defining macrophage heterogeneity in chronic intracellular bacterial infection in an unbiased manner is critical for understanding how macrophage functional diversity contributes to restricting microbes and limiting immunopathology, yet enables the pathogens to persist at low levels within granulomas and infected tissues for long periods of time. Here, we performed scRNA-seq to determine the functional diversity that underlies the capacity of macrophages to permit or limit bacterial persistence in the spleens of mice that have been chronically infected with *S*Tm for one month. We identified determinants of macrophage heterogeneity and described populations of distinct phenotypes, functional programming, and spatial localization. Using the *ΔsteE S*Tm mutant, which has a defect in counteracting host signals to skew macrophage phenotypes and maintain tissue persistence, we delineated macrophage phenotypes that contribute to controlling the infection. We found that angiotensin converting enzyme (ACE) expression defines a splenic macrophage population that localizes to granulomas and the abundance of the ACE^+^ macrophage niche anticorrelates with tissue bacterial burden. ACE^+^ granuloma macrophages rarely harbor persistent *S*Tm and granuloma macrophages with intracellular *S*Tm are much less likely to express ACE, indicating that ACE^+^macrophages are a non-permissive phenotype. Unbiased transcriptomics analysis revealed a differential enrichment of genes involved in pathogen response programming such as TNF signaling, type-I interferon response, and fatty acid metabolism in *Ace^+^* macrophages. Disruption of pathogen control by neutralizing TNF preferentially depletes ACE^+^ macrophages in infected spleens, compared to iNOS^+^ macrophages, which are a cellular niche for *S*Tm within granulomas. Collectively, our data suggest a model in which a balance between non-permissive ACE^+^ macrophages and *S*Tm macrophage niches in infected spleens defines controlled infection. Our study thus reveals ACE^*+*^ macrophages as a functionally distinct macrophage phenotype that could be targeted to limit intracellular bacteria persistence in infected tissues.

## Results

### The spectrum of splenic macrophages, monocytes, and their precursors during persistent *S*Tm infection

Single-cell RNA sequencing (scRNA-seq) is a powerful approach for identifying cellular phenotypes and their functional roles in an unbiased fashion(38, 39). We used scRNA-seq to comprehensively define macrophage heterogeneity and functional diversity in the spleens of 129×1/SvJ mice that were infected with fully virulent *S*Tm for 1 month, a time point at which the bacterial levels have been controlled and persistent infection has been established (4, 14). The SPI2 T3SS effector SteE polarizes macrophages to a more permissive state and thereby promotes *S*Tm persistence(14, 26, 28). As a result, mice infected with *ΔsteE S*Tm mutant have approximately 10-fold less bacteria in the spleens by 1 month post-inoculation, compared to WT *S*Tm-infected mice(14, 40). We performed scRNA-seq on splenocytes from both WT STm and *ΔsteE S*Tm-infected mice to gain insights into how macrophage states and phenotypes control bacterial persistence (Fig. S1A).

We previously showed that splenic *S*Tm granulomas are densely populated by CD11b^+^CD11c^+^Ly6C^+^ mononuclear phagocytes that express high levels of the phagocytosis receptor CD64, a marker universally expressed in macrophages across mouse tissues(14, 41). These granuloma macrophages also have high levels of MHCII and F4/80 expression consistent with activated macrophages(14) (Fig. S1B). The CD11b^+^CD11c^+^Ly6C^+^ granuloma macrophages (hereafter referred to as granuloma macrophages) are rare in the uninfected spleens but expand by 100-fold to constitute approximately 1% of total cells in the infected spleens by 1 month post-inoculation(14). Thus, to comprehensively define the phenotypes and functional features of these granuloma macrophages, their precursors, and other types of macrophages in *S*Tm-infected spleens, we devised a permissive fluorescence activated cell sorting (FACS)-enrichment strategy that simultaneously enriches for granuloma macrophages and captures other splenocytes for droplet-based scRNA-seq using 10X Genomics platform (Fig. S1A-B) (Methods). We performed two independent experiments, each with two WT *S*Tm and two *ΔsteE S*Tm-infected mice. We captured and sequenced a total of 40,281 cells. We filtered cells with low quality by removing those that had < 1000 unique molecular identifier (UMI) counts, < 500 detected genes, and > 5% reads of mitochondrial origin, resulting in 22,512 cells that passed quality controls (Fig. S1D-G). We detected on average 2,714 genes per cell and observed no substantial differences between individual experiments (Fig. S1D-G). We combined the filtered samples for downstream analysis.

To resolve cellular heterogeneity in the dataset, we applied an unsupervised, soft feature-learning method, SAM(42), to learn a cell-to-cell similarity matrix based on the intrinsic variation of gene expression between cells. Based on the resulting graph, we then inferred the cell types by clustering cells that share similar gene expression profiles (Fig. S2A). We annotated the cell type and designated the immune cell types of our scRNA-seq dataset by referencing the scRNA-seq PanglaoDB database(43). We captured all the major splenic immune cell types, as expected by our permissive FACS-enrichment strategy (Fig. 1A-C). Importantly, we found that each cell cluster contains a mixture of cells from different individual mice, suggesting that experimental batch effect is not a dominant source of variation in our dataset (Fig. S2B). Macrophages and monocytes, along with dendritic cells, are tissue sentinels that make up the mononuclear phagocyte system(44, 45). Tissue macrophages have dual developmental origins. In all mammalian tissues, a fraction of these cells originates from embryonic precursors and others share a direct lineage with monocytes(30, 45). Notably, mononuclear phagocytes (MNPs) that are not dendritic cells constitute over 37% of the total cell population (Fig. 1A-B), indicating that macrophages, monocytes, and their precursors are highly enriched in our dataset.

**Figure 1:**
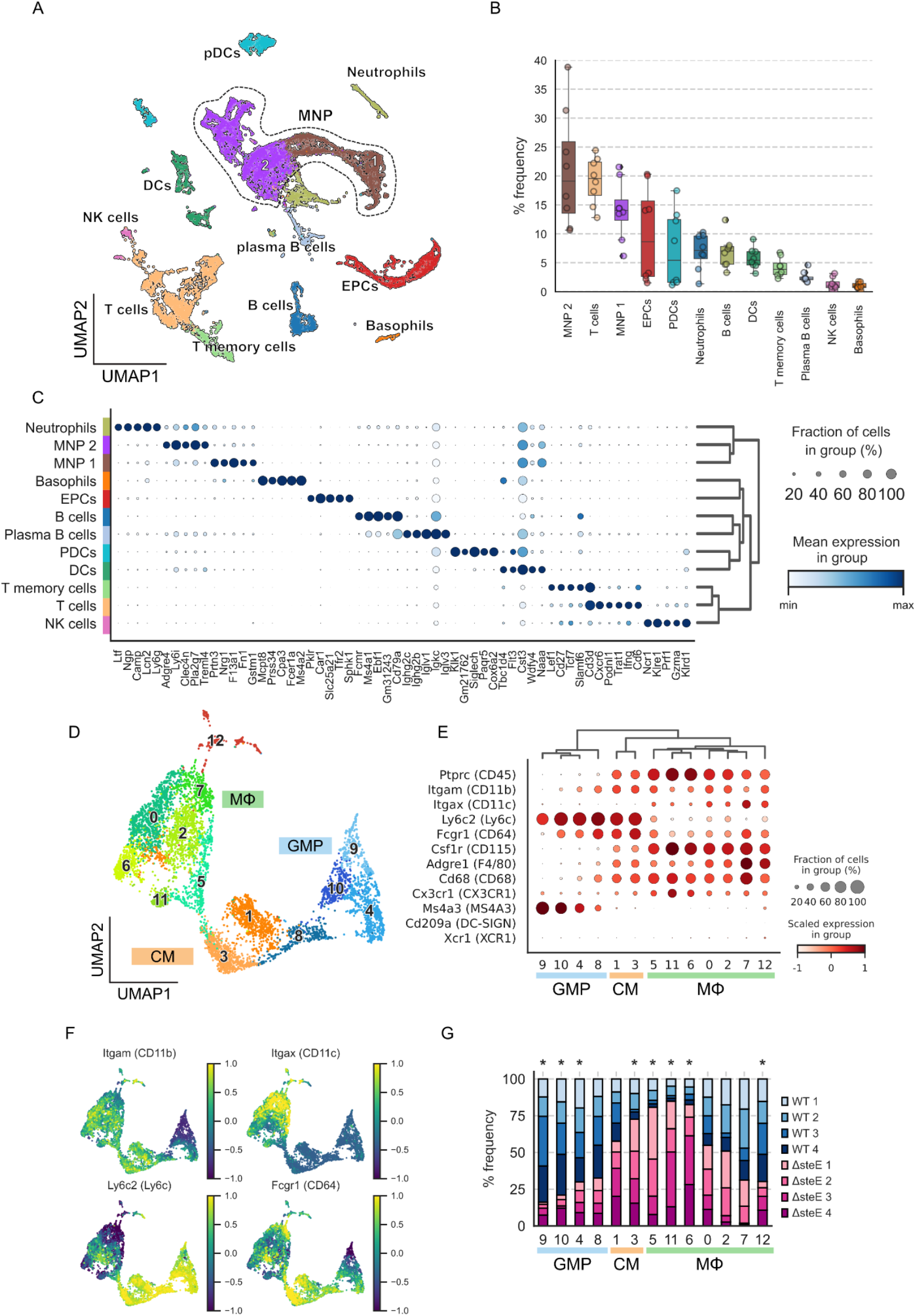
The spectrum of splenic macrophages, monocytes, and their precursors during persistent *S*Tm infection. Splenocytes from mice infected with either WT *S*Tm or *ΔsteE S*Tm for 4 weeks were permissively FACS-sorted to simultaneously enrich for macrophages and capture all immune cell types for scRNA-sequencing. **(A)** UMAP projection of sequenced cells from both WT *S*Tm and *ΔsteE S*Tm-infected spleens). **(B)** Percent frequency for each immune cell type. **(C)** Top 5 differentially expressed genes in each immune cell population. Radius and color intensity of the dots reflect detection rate and mean expression for each gene, respectively. **(D)** UMAP projection of mononuclear phagocytes (MNPs), excluding dendritic cells and neutrophils, colored by the assigned subpopulations: granulocyte mononuclear progenitors (GMP), classical monocytes (CM), and macrophages (MΦ). **(E)** Dotplot showing expression levels of myeloid marker genes on GMP, CM, or MΦ clusters. (F) Expression levels of *Itgam* (CD11b), *Itgax* (CD11c), *Ly6c2* (Ly6C), and *Fcgr1* (CD64) previously shown to express on macrophages that densely populate *S*Tm granulomas (Figure S1B). **(G)** Differential representation test for GMP/CM/MΦ clusters in WT *S*Tm and *ΔsteE S*Tm infected animals. Asterisk above the bar indicates a greater than 2-fold difference in representation ratio and statistical significance in association with bacterial strain based on differential representation test (FDR < 0.05).

Next, we performed subclustering of the MNP groups (demarcated by dotted boundary in Fig. 1A) to investigate the spectrum and heterogeneity of macrophages, monocytes, and their precursors more in depth. Myeloid cells are highly heterogeneous and exhibit overlapping transcriptional landscapes(46). Among the MNP clusters, we identified multiple distinct populations of macrophages (MΦ), classical monocytes (CM), and granulocyte-monocyte progenitors (GMP), which gives rise to monocytes and monocyte-derived macrophages(47) (Fig. 1D). We validated the sub-cluster identities by performing hierarchical clustering on a panel of myeloid lineage marker and functional genes (Fig. 1E). Notably the GMP clusters (GMP 9, 10, 4, and 8) express high levels of the GMP marker *Ms4a3*(48). Compared to the GMP and CM clusters (CM 1 and 3), the seven distinct macrophage clusters (MΦ 5, 11, 6, 0, 2, 7, and 12) express intermediate levels of *Ly6c2*. In addition, they co-express varying levels of *Itgam* (CD11b), *Itgax* (CD11c), *Fcgr1* (CD64), *Adgre1* (F4/80), and Csf1r (CD115) (Fig. 1E-F), indicating that they encompass the *S*Tm granuloma macrophages we previously identified(14). By contrast, the MΦ clusters express very low levels of *Cd209a* and *Xcr1*, which are known to be more highly expressed on dendritic cells.

To determine the impact of the SPI2 T3SS effector SteE on the functional heterogeneity of macrophages and macrophage precursors, we measured differential representation of GMP, CM, and MΦ phenotypes in WT STm-infected, compared to *ΔsteE S*Tm-infected samples (Fig. 1G) (Methods). We found that GMP clusters 4, 9, 10 and MΦ 12 are significantly more abundant in WT *S*Tm-infected spleens (FDR < 0.05. In contrast, CM cluster 3 and MΦ clusters 5, 6, and 11 are significantly more enriched in *ΔsteE S*Tm compared to WT *S*Tm infection. These findings suggest the SteE effector activity markedly altered the composition and functional diversities of macrophages, monocytes, and their precursors in infected tissues that may contribute to bacterial persistence and control of infection.

### Delineating distinct macrophage functional programming and phenotypes in infected spleens

Next, we used our single-cell transcriptomics analysis to determine the heterogeneity of macrophage populations with distinct functional states and phenotypes in the infected spleens. We performed differentially expressed gene (DGE) analysis and identified the most enriched gene sets for each of the GMP, CM, and MΦ clusters (Fig. 2A). This analysis demonstrates that these cell clusters are differentiated by distinct expression patterns of genes involved in macrophage functions and responses, including TNF signaling, cell death, type-I interferon, and complement activation (Fig. 2B). These macrophage functional activities are crucial for host immune response against bacterial pathogens(49–51). Innate cellular responses are typically thought to be immediate and early host immune response against pathogens. Remarkably, even at 1 month post-inoculation when *S*Tm infection is fully established in the spleens and the bacterial level has been controlled(14), we observed significant heterogeneity in macrophage functional responses, suggesting that tissue macrophage immunity is highly heterogenous during persistent infection *in vivo* and involves macrophages with varying antibacterial capacity.

**Figure 2:**
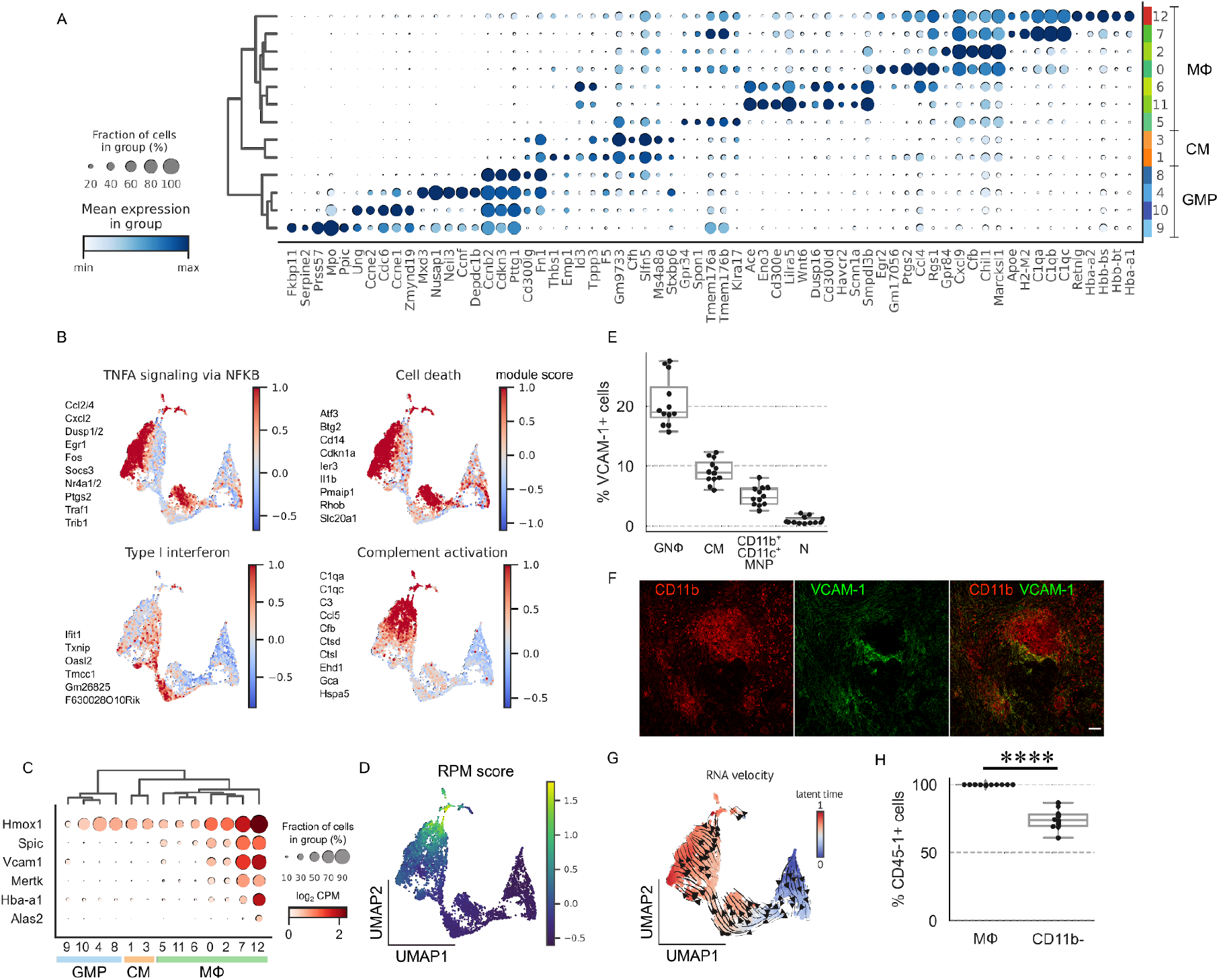
Delineating distinct macrophage functional programming and phenotypes in infected spleens. **(A)** Top 5 differentially expressed genes for GMP, CM, and MΦ clusters. Radius and color intensity of the dots reflect detection rate and mean expression for each gene, respectively. **(B)** Ensemble expression score of immune response gene sets using top differentially expressed genes (see Methods). Members of the gene sets are shown next to the plots. **(C)** Dotplot showing expression levels of red pulp macrophage (RPM) marker genes. **(D)** Ensemble expression score of RPM marker genes (*Hmox1, Spic, Vcam1, Mertk, Hba-a1, Alas2*).**(E)** Frequencies of VCAM-1 expression in different myeloid populations in WT *S*Tm-infected spleens at 1-month post-inoculation by flow cytometry. Granuloma macrophage (GNΦ), classical monocyte (CM), CD11b^+^CD11c^+^ mononuclear phagocyte (CD11b^+^CD11c^+^MNP), neutrophil (N). **(F)** Confocal imaging of splenic granuloma macrophages stained for CD11b (red) and VCAM-1 (green). **(G)** UMAP projection of GMP, CM, and MΦ clusters with predicted RNA velocity vector field overlaid on the top hints at developmental transition from progenitor state to VCAM-1^+^ macrophages. **(H)** Percent frequencies of CD45-1^+^ cells among VCAM-1 ^+^granuloma macrophages and CD11b^−^ cells in CD45.1^−^ recipient chimeric mice re-constituted with CD45.1^+^ bone marrow. **E**, **H**. Dot: individual mice. Significance calculated using a two-tailed Mann-Whitney test. **** p <0.0001.

In our analysis of the macrophage clusters, we discovered two clusters (MΦ 7 and 12) with uniquely high *Vcam1* expression (Fig. 2C). In the spleen, red pulp macrophages (RPM), which are thought to originate from embryonic precursors, had been shown to express VCAM1(52, 53). RPM development is dependent on the transcriptional factor SPIC and these macrophages play a role in heme metabolism through phagocytosing spent red blood cells(52, 53). To delineate the *Vcam1*-expressing MΦ 7 and 12 clusters, we examined the expression of RPM marker genes, including *Spic, Hmox1, Vcam1, Mertk, Hba-a1*, and *Alas2* (Fig. 2C). We found that these genes are highly expressed in both MΦ clusters 7 and 12, except Alas2, which is detected only in MΦ cluster 12. We calculated an ensemble score (see Methods) based on the expression of RPM marker genes and identified MΦ cluster 12 as RPMs (Fig. 2D). As shown in (Fig. 1E - F), *Vcam1^+^* MΦ cluster 7 also co-expresses CD11b (*Itgam*), CD11c (*Itgax*), Ly6C (*Ly6c2*), and CD64 (*Fcgr1*), suggesting that they may be macrophages that contribute to the formation of splenic granulomas during *S*Tm infection. To investigate this possibility, we performed flow cytometry analysis and found that VCAM1 is expressed on approximately 20% of granuloma macrophages, while other cell types of myeloid lineages have lower expression (Fig. S1B and 2E). By performing immunostaining on the *S*Tm-infected spleens, we found that VCAM1^+^ granuloma macrophages are spatially localized to the periphery of granulomas (Fig. 2F).

The developmental origins of macrophages that organize into granulomas in intracellular bacteria-infected tissues are still not well defined(54). Prior studies showed that under steady state, bone marrow-derived monocyte precursors give rise to a population of VCAM1^+^ splenic macrophages that regulate heme metabolism in the spleens(55). To test if the VCAM1^+^ granuloma macrophages in *S*Tm-infected spleens originate from a bone marrow origin, as opposed to embryonic precursors, we first performed RNA velocity analysis to gain insight into their development. Recent advances in scRNA-seq analysis have enabled the unbiased quantification of transcriptional trajectory based on kinetic modeling of mRNA splicing(56). To determine the transcriptional dynamics of *Vcam1*-expressing macrophage clusters, we applied RNA velocity to our scRNA-seq dataset. We determined an RNA-velocity trajectory that is concordant with monocyte-macrophage transition and suggests that the *Vcam1^+^* MΦ 7 cluster arises from bone-marrow derived GMPs (Fig. 2G). To validate this relationship, we generated bone-marrow chimera by replacing the hematopoietic compartment in lethally irradiated CD45.2^+^129×1/SvJ recipient mice with bone marrow cells from CD45.1^+^ 129×1/SvJ donor mice. Fully reconstituted chimeric mice were then infected with WT *S*Tm and analyzed at 1 month post-inoculation. We found that 100% of the VCAM1^+^ granuloma macrophages were derived from CD45.1^+^ donors, demonstrating that they originated from a bone marrow source. By contrast, a lower fraction of the CD11b-cells, which include radiation-resistant cells such as memory T cells, were derived from CD45.1^+^ donors (Fig. 2H). Collectively, these data demonstrate that our permissive-FACS enrichment approach and single-cell transcriptomics identify a wide spectrum of tissue macrophages with distinct functional states, phenotypes, and ontogenetic development in the spleen during persistent *S*Tm infection.

### Identifying macrophage phenotypes associated with limiting infection

To further investigate the functional diversity and identify macrophage phenotypes that contribute to controlling persistent *S*Tm infection in the spleens, we sub-clustered the MNPs in Figure 1D without the GMPs, which have high expression of cell-cycle related genes indicating proliferative activities (Fig. S3A-B). We identified 20 sub-clusters (Fig. 3A). Among these cells, clusters 18, 0, 3, 2, 6, 14 exhibit transcriptional signatures consistent with classical monocytes; whereas, the remaining clusters were identified as macrophages by their patterns of myeloid lineage and functional marker expression (Fig. 3B). We then performed DGE analysis for each macrophage cluster and designated MΦ populations based on the top expressing genes (Fig. 3C-D). Analysis of the top most defining genes (FDR < 0.05, expressed in > 10% of the cluster, log_2_ fold-change > 1) for each annotated cluster reveals different macrophage phenotypes and functional states (Fig. 3E). Among these, we identified RPMs (clusters 15-17) and *Vcam1^+^* granuloma macrophages (cluster 10), as characterized in Figure 2. We identified additional functional heterogeneity in the macrophage subpopulations. Cluster 12 defines a macrophage population that expresses *Nos2*, which encodes the inducible nitric oxide synthase enzyme (iNOS), that had been previously shown to be a distinct granuloma macrophage phenotype in *Mtb*-infected lungs of non-human primates and *S*Tm-infected spleens of mice(13, 14, 31). *Trem2^+^* macrophages (cluster 9), which were recently identified as a distinct macrophage subset in human *Mycobacterium leprae* granulomas, are also present in our dataset(57). *Egr^+^* macrophages (clusters 8 + 13) express high levels of Hbegf, Egr1, and Egr2, which are early activating transcriptional factors in macrophages(58). Our analysis also identified *Vcan* as a highly enriched gene in the monocyte clusters (Fig. 3C & E), which was shown to be a universal marker for monocytes across different human tissues based on scRNA-seq(45). Macrophage clusters 1 and 7 express high levels of Gpr84 and Celf4, respectively, though the expressions of these genes are relatively diffuse across several macrophage subpopulations (Fig. 3D - E). Intriguingly, we identified clusters 4 and 11 as macrophages that express high levels of angiotensin converting enzyme (Ace). ACE is a zinc-containing dipeptidyl carboxypeptidase that generates bioactive peptides regulating blood pressure, cardiovascular physiology, and inflammation(59). The role of ACE^+^ macrophages during persistent intracellular bacterial infections has not been defined.

**Figure 3:**
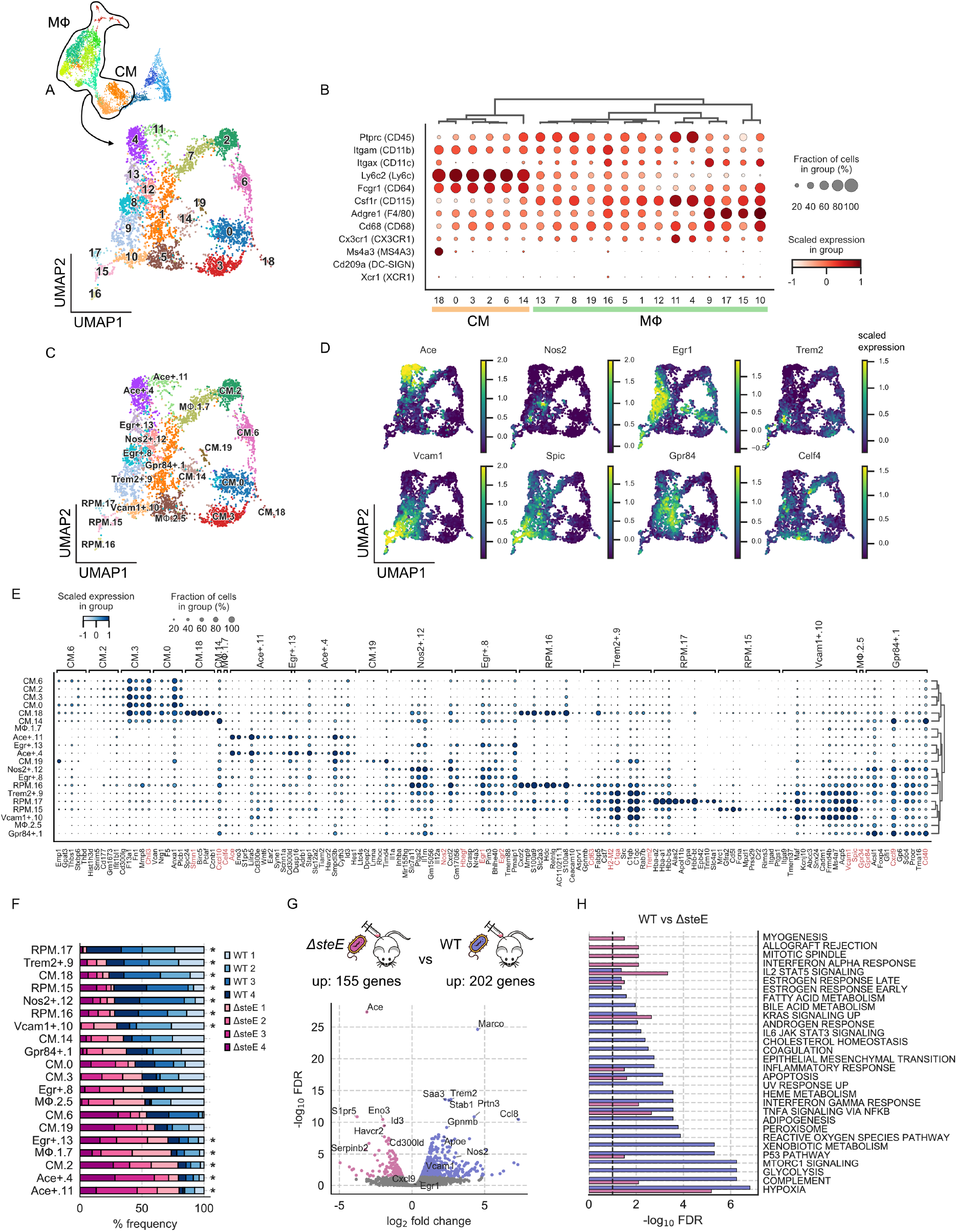
Identifying macrophage phenotypes associated with limiting infection. **(A)** UMAP projection of monocyte and macrophage subsets from WT *S*Tm and *ΔsteE S*Tm infected mice. Cells are colored by Leiden cluster assignment. **(B)** Expression levels and frequencies of myeloid cell marker genes. **(C)** UMAP projection of monocyte and macrophages subsets from WT *S*Tm and *ΔsteE S*Tm-infected mice. Cells are colored by cell state assignment. **(D)** Expression levels and frequencies of enriched marker genes of macrophage clusters: *Ace, Nos2, Egr1, Trem2, Vcam1, Spic, Gpr84, Celf4*. **(E)** Expression levels and frequencies of the top 5 representative differentially expressed genes (DEGs) from each cell state (FDR < 0.05, expressed in > 10% of cluster, log_2_ fold-change > 0.5). A select panel of genes that define the functional state is highlighted in red. **(F)** Differential representation test for monocyte and macrophage states in WT *S*Tm and *ΔsteE S*Tm-infected animals. Asterisk next to the bar indicates a greater than 2-fold difference in common odds ratio and statistical significance (<0.05 FDR) between the corresponding mice replicate (FDR < 0.05). **(G)**Volcano plot showing differentially gene expression analysis between combined macrophage populations with differential abundances (marked with asterisks in Figure **3F**) in WT *S*Tm and *ΔsteE S*Tm-infected spleens. −log_10_ FDR < 0.05 and log_2_ fold change > 0.5. Negative binomial test. **(H)** Gene sets over-representation analysis of the genes that are differentially up-regulated in the select macrophage populations (marked with asterisks in Figure 3F) of WT *S*Tm and *ΔsteE S*Tm-infected. Genes with FDR < 0.05, log_2_ fold-changes > 0.5, and > 10% expression in the corresponding group are selected as input for analysis. Bar width reflects the −log_10_ FDR.

The interactions between host signals and bacterial factors shape the balance between antibacterial and bacteria-permissive states in granuloma macrophages during infection(14, 22, 26). This balance affects bacterial control and infection outcome. The *ΔsteE S*Tm mutant has a defect in polarizing macrophages toward a permissive state, leading to reduced bacterial tissue persistence and more rapid control of the infection(14, 26, 28). To identify macrophage phenotypes that may contribute to limiting bacterial persistence and controlling infection, we compared the abundances of our scRNA-seq macrophage phenotypes in WT STm and *DsteE S*Tm-infected spleens. We performed a differential representation test to determine relative enrichment of monocytes and macrophages. Remarkably, we found that the majority of frequency variation is in the macrophage subpopulations (Fig. 3F). Our analysis showed that *DsteE S*Tm infection led to lower frequency of RPM, *Nos2*^+^, *Vcam-1*^+^, and *Trem2*^+^ macrophages, compared to WT *S*Tm infection. In contrast, the *Ace*^+^ (clusters 4 and 11), *Celf4* (cluster 7), and *Egr*^+^ (cluster 13) subpopulations are more abundant in *ΔsteE S*Tm-infected spleens, which have higher level of bacterial clearance.

To gain insights into the impact of bacterial effector SteE activity on macrophage functional pathways, we focused on the macrophage populations that exhibited differential representation between WT *S*Tm and *ΔsteE S*Tm infected animals (annotated with asterisks Fig. 3F). Combining these populations into a WT *S*Tm and a *ΔsteE S*Tm-infection group, we performed DEG analysis to determine the impact of SteE activity on macrophage transcriptional programming (Fig. 3G). We identified 155 genes that are significantly upregulated (FDR < 0.05 and log2 fold change > 0.5) in the macrophages from *DsteE S*Tm-infected animals. The top DEGs are enriched in *Ace+* macrophages, including *Ace*, *Eno3, S1pr5*, and *Id3*, consistent with our representation analysis that *Ace*^+^ macrophages were significantly enriched in *ΔsteE S*Tm-infected spleens. In contrast, 202 genes are significantly upregulated in the macrophages from WT *S*Tm-infected animals, many of which are genes enriched in RPMs and *Nos2*^+^ macrophages. We then quantified the enrichment of functional pathways by performing gene sets over representation (GSOA) analysis on the DEGs (see Methods). Despite robustly expressing genes involved in antibacterial responses such as TNF and IFNγ signaling to a similar extent, splenic macrophages from the WT *S*Tm-infected mice are markedly more enriched in genes involved in complement activation, peroxisome, reactive oxygen species, glycolysis, heme metabolism, and adipogenesis, demonstrating remarkable SteE-driven effects on a wide range of macrophage immune and metabolic activities (Fig. 3H). Collectively these results suggest that *Ace*+ macrophages are one of the phenotypes that drive the functional transcriptional differences between macrophages in WT STm and *ΔsteE S*Tm spleens and their abundance is linked to infection containment.

### Splenic ACE^+^ macrophages expand during infection and contribute to granuloma formation

Our transcriptomics analyses suggest *Ace*^+^ macrophages are a phenotypically and functionally distinct population of macrophages in infected tissues during persistent *S*Tm infection. Intriguingly, ACE expression had been observed in human *Mtb* and sarcoidosis granulomas, suggesting that ACE^+^ macrophages may be commonly involved in granulomatous response across different persistent intracellular bacterial infections and pathophysiologic settings(60, 61). However, whether ACE^+^ macrophages are a functionally distinct population of granuloma macrophages and what role they play during persistent intracellular bacterial infections remain unknown. To delineate *Ace*-expressing macrophages, we first identify them in infected tissues and determine their tissue dynamics during persistent *S*Tm infection in the spleen. We performed flow cytometry analysis and found that among cells of granuloma macrophage phenotype (Fig. S1B), 10-20% express ACE, indicating that ACE^+^ macrophages are a subset of granuloma macrophages. The frequencies of ACE^+^ cells are significantly lower among CM and CD11b^+^CD11c^+^ MNPs and are marginally above background staining among neutrophils (Figure 4A, S4A-C). ACE^+^ macrophages are rare in uninfected spleens but by 2 weeks post-inoculation, their frequencies expand by over 10-fold in the infected spleens (Fig. 4B), which have significantly higher total cell numbers compared to uninfected spleens(14). By contrast, the frequency of splenic CD11b^+^CD11c^+^ MNP remains relatively constant over the course of infection. To determine the spatial localization of ACE^+^ macrophages within *S*Tm splenic granulomas, we performed confocal microscopy and found that they are distributed throughout granulomas (Fig. 4C). These data demonstrate that ACE^+^ macrophages are a distinct tissue macrophage population that expands in response to *S*Tm infection and contributes to splenic *S*Tm granuloma formation.

**Figure 4:**
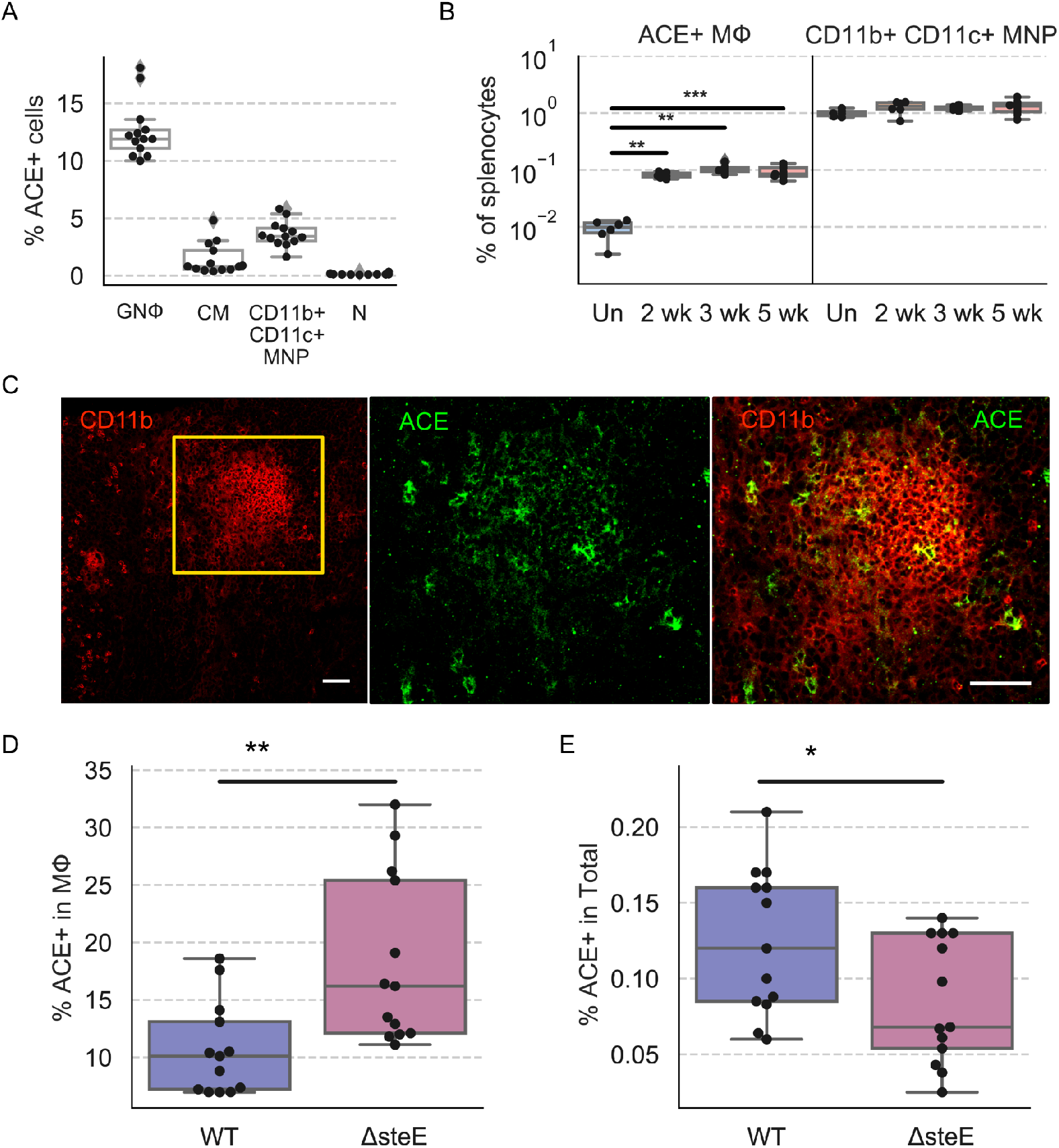
Splenic ACE^+^ macrophages expand during infection and contribute to granuloma formation. **(A)** Mice were infected with WT *S*Tm and analyzed at 1-month post-inoculation. Percent frequencies of ACE^+^ cells among granuloma macrophages (GNΦ), classical monocytes (CM), CD11b^+^CD11c^+^ mononuclear phagocytes (MNP), and neutrophils (N) by flow cytometry. **(B)** Percent frequency of ACE^+^ granuloma macrophages (left) and CD11b^+^CD11c^+^ mononuclear phagocytes (MNPs) (right) in total splenocytes at indicated time points post-inoculation. Un: uninfected. **(C)** Immuno-fluorescence staining *S*Tm granuloma macrophages for CD11b (red) and ACE (green). Orange square on the left panel indicates the zoomed region shown in the two right panels. **(D-E)** Mice were infected with either WT *S*Tm or *ΔsteE S*Tm and analyzed at 1-month post-inoculation by flow cytometry. **(D)** Percent frequencies of ACE^+^ cells among granuloma macrophages. **(E)** Percent frequencies of ACE^+^ granuloma macrophages among total splenocytes. **A**, **B**, **D**, **E**. Dot: individual mice. Significance calculated using a two-tailed Mann-Whitney test. * p < 0.05, ** p < 0.01, *** p < 0.001.

We next investigated whether the ACE^+^ granuloma macrophage phenotype is associated with controlling *S*Tm infection in the spleens by comparing their tissue levels in WT and *DsteE S*Tm-infected mice, which have reduced bacterial persistence(14). We found that by one month post-inoculation, ACE^+^ cells were significantly more abundant among granuloma macrophages in *ΔsteE S*Tm-infected spleens, compared to WT *S*Tm-infected spleens (Fig. 4D). The percent frequencies of ACE^+^ granuloma macrophages among total splenocytes were slightly lower in the *ΔsteE S*Tm-infected spleens (Fig. 4E), which weigh approximately 5-fold less and have reduced total cell numbers, compared to WT *S*Tm-infected spleens(14). These findings suggest that an increased abundance of ACE^+^ cells among granuloma macrophages may contribute to reducing *S*Tm persistence in the infected spleens.

### ACE^+^ granuloma macrophages are a non-permissive cellular niche for *S*Tm

The increased abundance of the ACE^+^ macrophage niche in the *ΔsteE S*Tm-infected spleens, which have markedly reduced tissue bacterial levels, led us to postulate that ACE^+^macrophages contribute to limiting *S*Tm persistence and infection. Thus, we determined the capacity for ACE^+^ macrophages to act as a cellular niche of intracellular *S*Tm by measuring the frequencies of *S*Tm-containing cells among these macrophages using flow cytometry. As expected with persistent *S*Tm infection in 129×1/SvJ mice, the pathogen is controlled at low chronic levels in the infected spleens by 1 month post-inoculation and approximately 0.2% of splenic granuloma macrophages contain intracellular bacteria at this stage of the infection(4, 14)(Fig. 5A). However, we found that splenic ACE^+^ granuloma macrophages are markedly less likely to harbor intracellular *S*Tm compared to ACE^−^ cells (Fig. 5A). Furthermore, *S*Tm-containing granuloma macrophages were approximately 5-fold more likely to have undetectable ACE expression, compared to *S*Tm-negative granuloma macrophages (Fig. 5B). These findings indicate that ACE^+^ granuloma macrophages are a non-permissive niche for *S*Tm.

**Figure 5:**
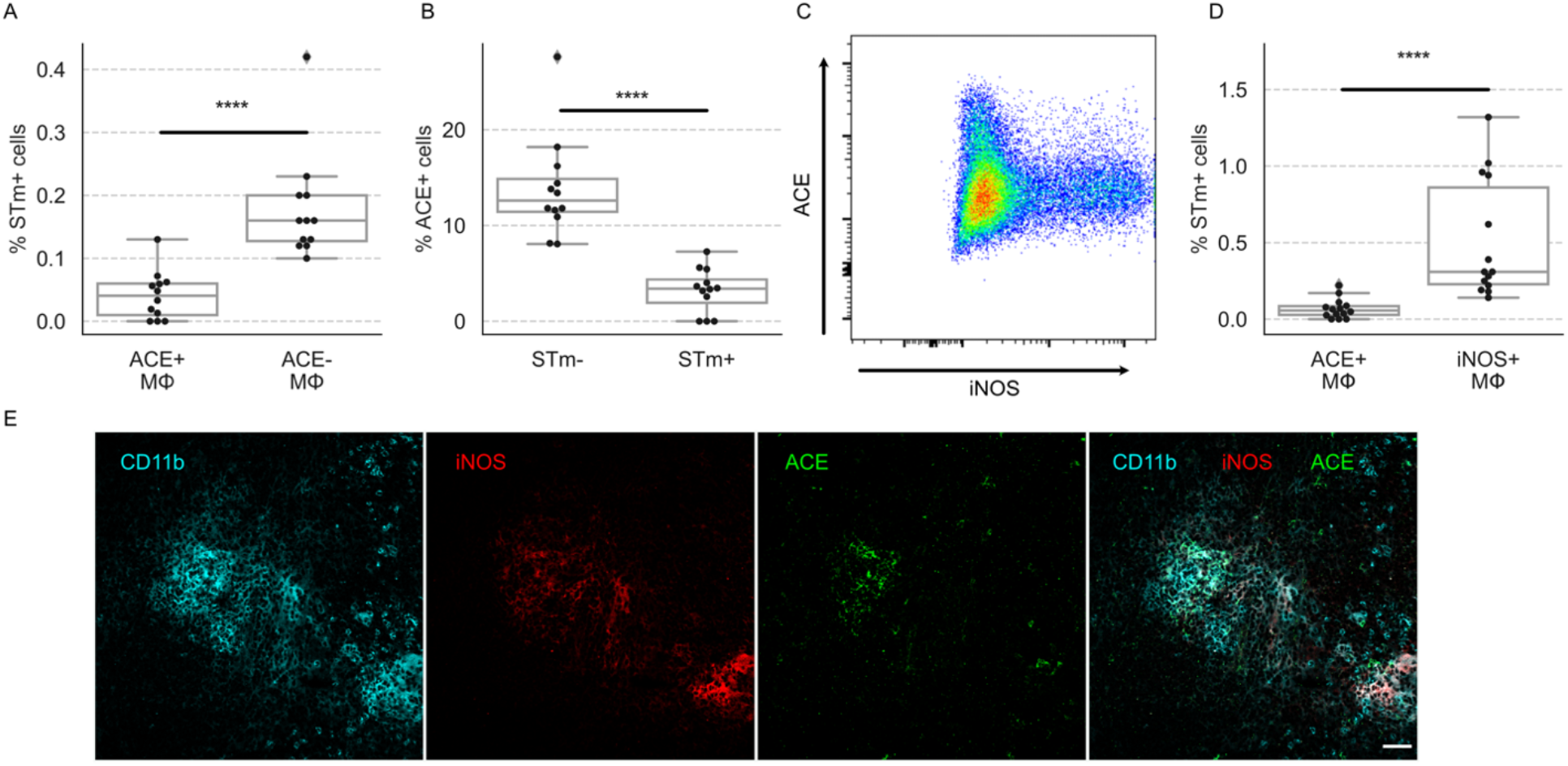
ACE^+^ granuloma macrophages are a non-permissive cellular niche for *S*Tm. Mice were infected with WT *S*Tm and analyzed at 1-month post-inoculation. **(A)** Percent frequencies of *S*Tm-infected cells among ACE^+^ and ACE^−^ granuloma macrophages by flow cytometry. **(B)** Percent frequencies of ACE^+^ cells among granuloma macrophages that are infected with *S*Tm. **(C)** ACE^+^ and iNOS^+^ granuloma macrophages form highly disparate populations. Splenocytes were analyzed by flow cytometry. Cells were first gated for granuloma macrophages as shown in Figure S1B, the plotted for ACE and iNOS expression as shown. **(D)** Percent frequencies of *S*Tm-infected cells among ACE^+^ and iNOS^+^ granuloma macrophages by flow cytometry. **(E)** ACE^+^ and iNOS^+^ macrophages have overlapping distribution within granulomas. Immuno-fluorescence staining granuloma macrophages for CD11b (cyan), iNOS (red), and ACE (green) in splenic granuloma macrophages. Merged channels are shown on the bottom right panel. **A**, **B**, **D**. Dot: individual mice. Significance calculated using a two-tailed Mann-Whitney test. **** p <0.0001.

Although iNOS expression has commonly been associated with proinflammatory, antibacterial macrophages, we and others have observed that *S*Tm can persist within iNOS^+^ splenic granuloma macrophages even at 1 to 2 months post-inoculation(13, 14)(Fig. S5A-C). To see if ACE^+^ and iNOS^+^ granuloma macrophages are phenotypically and functionally distinct populations, we analyzed splenocytes from infected mice for expression of these markers using flow cytometry. We found that ACE^+^ and iNOS^+^ cells form two largely non-overlapping granuloma macrophage subsets (Fig. 5C). Remarkably, the frequency of iNOS^+^ granuloma macrophages infected with *S*Tm was 8-fold higher than ACE^+^ granuloma macrophages (Fig. 5D). We next examined the spatial localization of ACE^+^ and iNOS^+^ macrophages to determine if ACE^+^ macrophages were excluded from iNOS^+^ centers of granulomas, which was previously thought to be an underlying factor that CXCL9/CXCL-10^+^ splenic macrophages are less likely to be infected with persisting *S*Tm(13). We found that ACE^+^ and iNOS^+^ granuloma macrophages have overlapping distribution within *S*Tm granulomas (Fig. 5E). Taken together, our data demonstrate that ACE^+^ granuloma macrophages are a distinct, non-permissive macrophage niche for intracellular *S*Tm in infected tissues.

We then sought to determine if ACE functionally controls the capacity of ACE^+^ macrophages to harbor intracellular *S*Tm. Transgenic mice (called Ace 10/10 mice) that overexpress *Ace* in myeloid cells due to ectopic placement of *Ace* under the *Csf1r* promoter have lower bacterial burdens during *MRSA* and *L. monocytogenes* infection at 3-5 days post-inoculation(62, 63). Peritoneal macrophages from *Ace* 10/10 mice exhibit an exaggerated proinflammatory response, with enhanced TNF, IL-6, and iNOS production, suggesting that ACE^+^ macrophages are more proinflammatory. In contrast, ACE-expressing human macrophages have been shown to have lower levels of proinflammatory cytokines such as TNF and IL-6(64). Furthermore, intracellular bacteria that can cause persistent infection, such as *S. enterica* and *B. henselae*, possess mechanisms to modulate macrophage responses and skew macrophage phenotypes(14, 26, 28, 29). Thus, the impact of ACE function on the capacity of macrophages to act as a cellular niche for intracellular bacterial persistence in infected tissues is unknown. To test if ACE expression is sufficient to alter macrophage permissiveness to *S*Tm, we expressed *Ace* in RAW264.7 macrophages, which have undetectable Ace expression at baseline or during *S*Tm infection, using lentiviral transduction *in vitro* (Figure S5D). Although transduced macrophages robustly expressed ACE, the levels of intracellular WT *S*Tm were not affected (Fig. S5D). We then sought to determine the impact of *Ace*-overexpression in myeloid cells on *S*Tm infection *in vivo*. We crossed ACE 10/10 mice, which are of C57BL/6 background that is highly susceptible to WT *S*Tm, with 129×1/SvJ mice, which are able to control *S*Tm infection(4). We used the mixed background ACE 10 F1 offspring to perform persistent infection (Fig. S5E). As expected, granuloma macrophages in the *S*Tm-infected spleens of ACE 10 F1 mice, which are heterozygous for ACE 10 transgene, expressed significantly higher ACE levels compared to control F1 mice (Fig. S5E-F). However, both groups of mice have similar splenic bacterial levels at 2 weeks post-inoculation (Fig. S5G). Together, these findings suggest ACE overexpression in macrophages alone is insufficient to impact *S*Tm persistence during *in vitro* and *in vivo* infection.

In addition, we tested to see if inhibition of ACE enzymatic activity affects *S*Tm tissue persistence by infecting 129×1/SvJ mice with WT *S*Tm for 1 month and then treating them either with saline control or the ACE inhibitor captopril, 150 mg/kg/day intraperitoneally, for 7 days. This dose of captopril was previously shown to abolish approximately 95% of splenic ACE enzymatic activity in mice infected with *Histoplasma capsulatum* and is 30-50 fold higher than the maximum daily captopril dose used in treating cardiovascular disease in humans(65). We observed no significant difference in splenic bacterial levels from captopril treatment during persistent *S*Tm infection *in vivo* (Fig. S5H). Collectively, our combined ACE overexpression and ACE enzymatic inhibition studies suggest that ACE is a defining marker of a non-permissive macrophage niche for *S*Tm and their capacity to act as a cellular niche for *S*Tm persistence is controlled by additional pathways beyond ACE.

### Disruption of pathogen control by TNF neutralization preferentially depletes ACE^+^ macrophages

Our data indicate that ACE expression defines a distinct granuloma macrophage niche that is non-permissive for *S*Tm and this macrophage population has strikingly disparate capacity to harbor intracellular *S*Tm compared to iNOS^+^ granuloma macrophages during persistent infection (Fig. 5A-D). Concordant with their phenotypic difference, unbiased pathway analyses revealed differential enrichment of genes involved in multiple pathogen response programming such as TNF signaling, type-I interferon response, fatty acid metabolism, and hypoxia between *Ace*^+^ macrophages and *Nos2*^+^ macrophages (Fig. S3C). Thus, to gain further insights into the cellular features and functional pathways underlying the phenotype of ACE^+^ macrophages and their regulations, we leveraged our transcriptomics data and compared the functional features of ACE^+^ macrophages with those of iNOS^+^ macrophages. We focused on cytokine and cytokine receptor expression, as cytokine signaling pathways, including IL4/IL-13, IL-10, IL-1, TNF, IL-18, and IL-6 are key determinants of macrophage immune and metabolic states, antibacterial functions, and capacity to permit intracellular bacterial persistence (14, 26, 28, 29, 33, 66, 67). We found that *Ace*^+^ and *Nos2*^+^ macrophages exhibit remarkably different cytokine and cytokine receptor gene expression patterns. While having similarly robust *Il1b* expression, the *Ace^+^*macrophages express significantly lower levels of *Tnf*, *Il6*, and *Il15ra*, which mediate pro-inflammatory, antibacterial immune responses (Fig. 6A). However, compared to *Nos2^+^*macrophages, *Ace^+^* macrophages also express markedly less *Il18bp* and *Il1rn*, both of which encode natural antagonists that limit the IL-18 and IL-1 pro-inflammatory effects, respectively(68, 69). *Ace*^+^ macrophages express similar levels of *Il4rα*, a commonly utilized marker for antiinflammatory, alternatively activated macrophages, but significantly higher levels of *Il10rα* compared to *Nos2*^+^ macrophages (Fig. 6A). In contrast, *Ace*^+^ macrophages also express higher levels of *Il6rα* and *Il17rα*, which transmit antimicrobial, pro-inflammatory signals. Collectively, our transcriptomics analyses demonstrate that *Ace*^+^ and *Nos2*^+^ macrophages have remarkably distinct transcriptional programming, with many contrasting cellular features that underlie their differential phenotypes and capacity to harbor *S*Tm and suggest they may be divergently regulated to control bacterial persistence in infected tissues.

**Figure 6:**
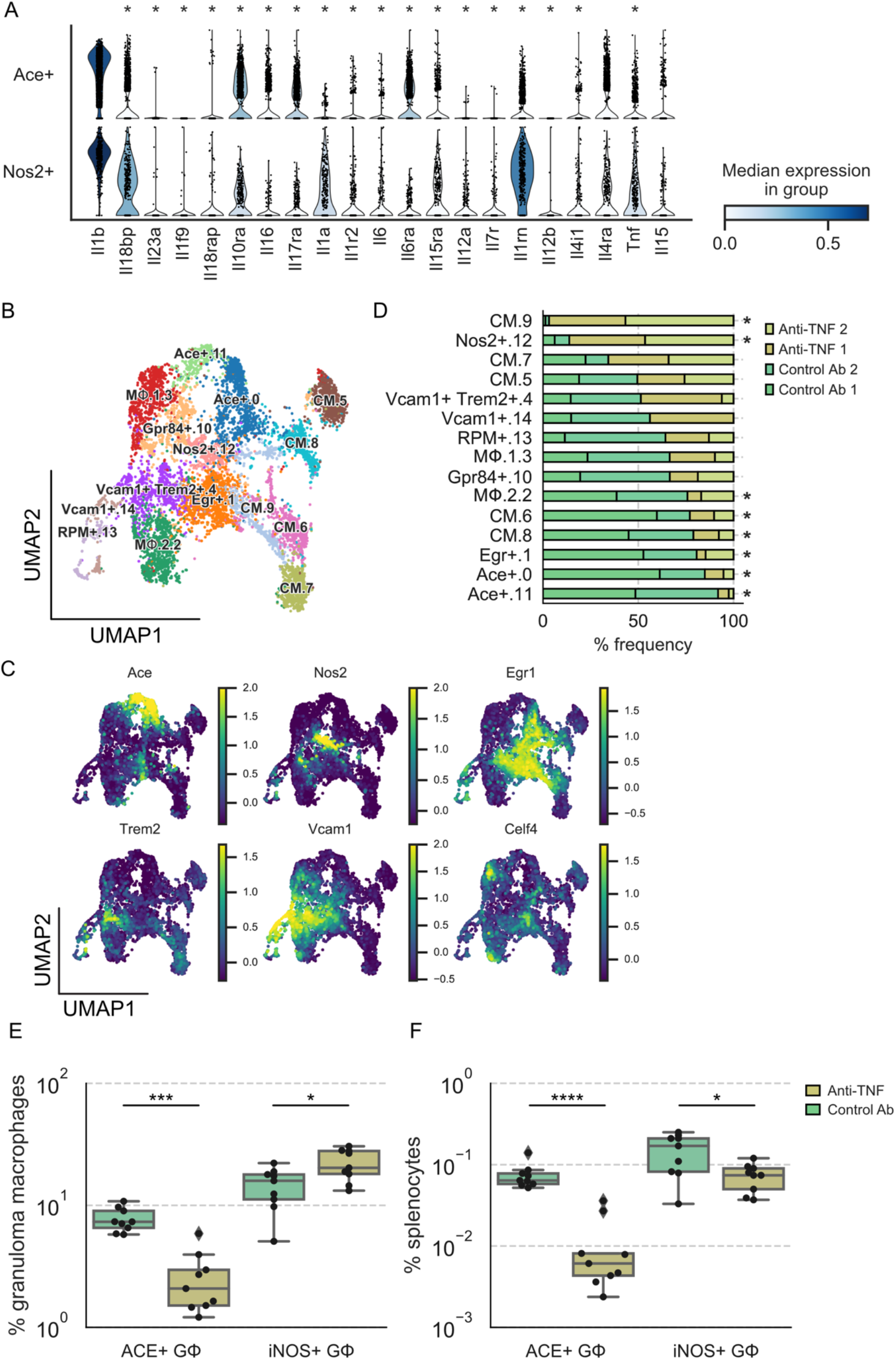
Disruption of pathogen control by TNF neutralization preferentially depletes ACE^+^ macrophages. **(A)** Violin plot highlighting cytokines that are differentially expressed between *Ace*^+^ and *Nos2*^+^ macrophages in WT *S*Tm or *ΔsteE S*Tm infected mice (see Figure **3A**) (FDR < 0.05). **(B)** Combined UMAP projection of scRNA-seq monocyte and macrophage subsets from mice that were treated with isotype control or anti-TNF antibody. Cells are colored by cell state assignment. **(C)**Expression levels of enriched marker genes of macrophage clusters: *Ace, Nos2, Egr1, Trem2, Vcam1, Celf4*. **(D)** Differential representation test for monocyte and macrophage clusters in WT *S*Tm-infected mice that have been treated with isotype control or anti-TNF antibody. Asterisk next to the bar indicates a greater than 2-fold difference in representation ratio and statistical significance between treatments based on a differential representation test (FDR < 0.05; see Methods). Mice were chronically infected with WT *S*Tm for 1 month, treated with isotype control or anti-TNF antibodies on day 0 and analyzed by flow cytometry or prepared for scRNA-seq on day 4. **(E)** Percent frequencies of ACE^+^ and iNOS^+^cells among splenic granuloma macrophages from animals treated with isotype control and anti-TNF antibody. **(F)** Percent frequencies of ACE^+^ and iNOS^+^ granuloma macrophages in total splenocytes from isotype control and anti-TNF treated animals. **A**, **F**. Dot: individual mice. Significance calculated using a two-tailed Mann-Whitney test. * p < 0.05, *** p < 0.001, **** p <0.0001.

Our data demonstrate that ACE^+^ granuloma macrophages are a non-permissive cellular niche for *S*Tm to persist within (Fig. 5A-B) and the abundance of this macrophage phenotype is significantly reduced in *ΔsteE S*Tm-infected spleens (Fig. 4D), which have reduced bacterial tissue persistence(14, 40). This led us to wonder if increased splenic *S*Tm persistence and loss of pathogen restriction may be associated with depletion of ACE^+^ granuloma macrophages. To probe the relationship between splenic ACE^+^ macrophage abundance and *S*Tm tissue bacterial levels, we disrupted pathogen control in infected mice by neutralizing TNF, which we had shown previously leads to increased *S*Tm splenic levels by 10-fold(14). The TNF neutralization effect was mechanistically linked to skewing macrophages toward a more bacteria-permissive state and the loss of pro-inflammatory macrophage phenotypes (14, 70, 71). Thus, we treated mice that had been infected with *S*Tm for 1 month with either control or anti-TNF neutralizing antibody. Since we observed striking differential enrichment of TNF signaling genes between *Ace*^+^ and *Nos2*^+^ macrophages (Fig. S3C), we examined both of these macrophage populations to see if TNF differentially affects *Ace*^+^ and *Nos2*^+^ macrophage abundance to control *S*Tm tissue persistence. Splenocytes were enriched for macrophages and monocytes using permissive FACS-enrichment strategy as described (see Methods). Samples from two control antibody-treated and two anti-TNF antibody-treated animals were subjected to scRNA-seq. We combined sequenced single-cell transcriptomes from the antibody-treated animals with the transcriptomes obtained from the WT *S*Tm and *ΔsteE S*Tm infection experiments (Fig. 3A) for a direct comparison of the same cell types. We observed that all macrophage and monocyte clusters, including the *Ace*^+^ and *Nos2*^+^ macrophage populations were comprised of cells from all animals across experiments (Fig. 6B-C, S7A), suggesting a lack of experiment-specific batch effect. As expected, differential representation test showed that *Ace*^+^ macrophages were more abundant in *ΔsteE S*Tm-infected spleens than in WT *S*Tm-infected spleens (Fig. S7B). Remarkably *Ace*^+^macrophages were more dramatically depleted in the spleens of animals treated with TNF neutralizing antibody, compared to animals treated with control antibody (Fig. 6D). In contrast, *Nos2*^+^ macrophages were significantly more abundant. To experimentally test if TNF neutralization disparately affects the *Ace*^+^ and *Nos2*^+^ macrophage niches in infected spleens, we performed flow cytometry analysis of *S*Tm infected mice that have been treated with either control or TNF neutralizing antibody. We found that among splenic granuloma macrophages, the percent frequencies of ACE^+^ cells were reduced by 4-fold in anti-TNF treated mice, compared to control antibody-treated mice. In contrast, the percent frequencies of iNOS^+^ cells increased slightly by 1.6 fold (Fig. 6E). Strikingly, the percent frequencies of ACE^+^ granuloma macrophages among total splenocytes were reduced by almost 10-fold in anti-TNF treated mice compared to control mice; whereas the percent frequencies of iNOS^+^ granuloma macrophages were reduced by only 2-fold (Fig. 6F). Taken together, our data suggest that TNF neutralization disrupts the control of *S*Tm persistence in infected spleens by preferentially depleting the non-permissive niche of ACE^+^ granuloma macrophages.

## Discussion

The tissue persistence of intracellular bacteria such as *Salmonella enterica, Brucella* species, and mycobacteria hinders treatment effectiveness and facilitates spread of infections(10, 11, 16). Despite robust innate and adaptive immune responses, in many individuals these pathogens can remain in infected tissues at low chronic levels for long periods of times and the hosts may not exhibit overt clinical symptoms. Central to this type of bacterial tissue persistence is the functional diversity and heterogeneity of macrophages and macrophage granulomas (12–14, 32, 35). In infected tissues, macrophages mediate critical antibacterial immune responses that contribute to pathogen eradication, resolution of inflammation, and tissue repair. Yet these crucial immune cells can also act as a cellular niche and form granulomas, which are immunological structures that function as a host mechanism to contain infection but within which intracellular bacteria are able to survive. Understanding of macrophage functional diversity requires delineation not only of macrophages within granulomas but also precursor cells that give rise to these and other macrophages in the infected tissue environment(32, 35, 37, 57). Here, we apply single-cell transcriptomics to investigate macrophage heterogeneity in *S*Tm-infected spleens to gain insight into how their heterogeneity and functional diversity contribute to controlling bacterial persistence and infection. Our permissive FACS-enrichment strategy not only enables significant enrichment of rare macrophage populations in infected tissues, such as the CD11b^+^CD11c^+^Ly6C^+^ granuloma macrophages, but also captures a full spectrum of splenocytes to facilitate comprehensive mapping of the tissue macrophage phenotypes during persistent infection (Fig. 1 and 2). In addition, we detected more than 2700 genes per cell on average, which provides substantial sequencing depth for delineating differences among macrophage populations. Indeed, our single-cell transcriptomics dataset captures red pulp macrophages, a known type of macrophages in the spleen. We also identify *Trem2*^+^ macrophages, which were recently identified as a macrophage subset in human *M. leprae* granulomas(57), as well as *Nos2*^+^ macrophages, a distinct macrophage phenotype in *M. tuberculosis* and *S. enterica* granulomas(13, 14, 31). Importantly, our single-cell transcriptomics enable delineation of macrophage phenotypes in granulomas and infected tissues during persistent intracellular bacterial infection that have not been defined, such as the bone marrow-derived VCAM-1 ^+^ granuloma macrophage population (Fig. 2) and ACE^+^ granuloma macrophage population (Fig. 3).

While ACE expression in *M. tuberculosis* granulomas had been well described, whether ACE^+^ macrophages are a distinct subset of macrophages in granulomas and what their functions are during persistent intracellular bacterial infection remain unknown. Our single-cell transcriptomics and functional characterization demonstrate that ACE expression specifies a macrophage population that has distinctive cellular features, functional properties, and TNF-regulation compared to other types of macrophages in *S*Tm granulomas and infected spleens (Fig. 4-6). The ACE^+^ granuloma macrophages described here are characterized by their expression of lineage markers CD11b, CD11c, and Ly6C, which is expressed at lower level compared to classical monocytes (Fig. S1B-C), suggesting that they may be monocyte-derived macrophages(72). It is now well recognized that in almost all tissues, some macrophages originate from blood-borne monocytic precursors recruited to the tissues during steady state or in the setting of inflammation; whereas, other macrophages arise from embryonic origin during development(30). Intriguingly, *M. tuberculosis* granulomas in various tissues in human and nonhuman primates are also populated with CD11b^+^CD11c^+^ macrophages(31, 73), raising the possibility that these macrophages might be analogous to the CD11b^+^CD11c^+^Ly6C^+^ granuloma macrophages we have described. Additionally, the ACE expression detected in human tuberculosis granulomas may reflect a functional equivalent ACE^+^ macrophage subset to the ACE^+^ granuloma macrophages that we report in this study. Furthermore, ACE expression has been observed in granulomas in the autoimmune disease sarcoidosis(60, 74). Collectively these reports and our present study suggest ACE^+^ macrophages might be involved in tissue granulomatous response across different types of tissues and diseases.

We found in this study that ACE^+^ macrophages in *S*Tm-infected spleens are less likely to harbor intracellular *S*Tm and conversely, *S*Tm-containing macrophages rarely have detectable ACE expression, indicating that ACE^+^ macrophages are non-permissive cellular niche for *S*Tm during persistent infection (Fig. 5A-D). By not harboring intracellular bacteria like other macrophages in granulomas and infected tissues, ACE^+^ macrophages may help limit the tissue persistence of *S*Tm. Our findings that the abundance of ACE^+^ macrophages is higher in *ΔsteE S*Tm-infected spleens, which have reduced bacterial persistence, but lower in spleens of TNF-neutralized animals, which have increased bacterial tissue levels and uncontrolled infection, suggest a model in which ACE^+^ macrophage abundance in infected tissues contributes to limiting bacterial persistence and infection. Prior studies using *Ace* transgenic mice showed that overexpression of ACE in *Csf1r*-expressing myeloid cells resulted in enhanced clearance of *MRSA* and *L. monocytogenes* from infected tissues at 3-5 days post-inoculation(63). In our persistent *S*Tm infection model, gain-of-function and loss-of-function manipulations of the ACE pathway bear no significant impact of bacterial persistence during *in vitro* or in *in vivo* infection (Fig. S5). We suspect that unlike *MRSA* and *L. monocytogenes*, vacuolar intracellular bacteria that cause persistent infection such as *S*Tm have a number of bacterial effector mechanisms to modulate macrophage responses and skew macrophage phenotypes(13, 14, 26, 28). Future studies involving selective deletion of *Ace* in tissue macrophages will further clarify on a potential impact of the *Ace* pathway on the non-permissiveness of ACE^+^ macrophages during persistent *S*Tm infection.

While ACE expression is a defining feature of a non-permissive macrophage niche, our data suggest additional ACE-independent pathways may influence the capacity of ACE^+^macrophages to harbor *S*Tm. By using ACE expression as a marker, we have been able to track and interrogate the differential cellular features and functional properties of ACE^+^ granuloma macrophages to gain insights into their functions and regulation. We found that ACE^+^ and iNOS^+^granuloma macrophages have vastly different capacities to harbor *S*Tm (Fig. 5). The non-permissive, ACE^+^ phenotype is likely a composite functional outcome of many cellular pathways that affect macrophage antibacterial activities and intracellular bacterial persistence. Examination of the cytokine signaling gene expressions showed that *Ace*^+^ macrophages in *S*Tm-infected tissues exhibit a pattern of cytokine and cytokine receptor expressions that does not neatly fit into either a pro-inflammatory or anti-inflammatory categorization(63, 64). Compared to *Nos2*^+^ macrophages, *Ace*^+^ macrophages express less *Tnf, Il1a, Il6*, and *Il15r* but more *Il6r* and *Il17r*, all of which are associated with pro-inflammatory signaling (Fig. 6). On the other hand, they also express higher levels of *Il10r*, which mediates an anti-inflammatory signal. Intriguingly, a recent study demonstrated that human macrophages lacking *Il10r* unexpectedly had reduced ability to restrict intracellular *S*Tm(75). Their findings suggest that macrophage *Il10r* expression is associated with a more bactericidal state with different cellular metabolic activities, including altered prostaglandin levels, that are unfavorable for intracellular *S*Tm persistence(75). In addition, we observed that *Ace*^+^ macrophages have markedly lower *Il18bp* and *Il1rn* expressions, compared to *Nos2*^+^ macrophages (Fig. 6). Deficiency of IL-18bp and IL-1Rn are thought to cause exaggerated pro-inflammatory responses in monocytes and macrophages and contribute to the development of inflammatory disorders such as Macrophage Activation Syndrome and autoimmune arthritis (68, 69). Corresponding with their differential cellular features, we found that TNF neutralization resulted in a preferential depletion of ACE^+^ macrophages (Fig. 6). TNF is a highly pleiotropic cytokine that has been linked to restraining the emergence of bacteria-permissive macrophage phenotypes and modulating macrophage cell death(14, 70, 71, 76). We speculate that the differential signaling state of ACE^+^ macrophages may result in disparate impacts on the fate of these cells upon TNF neutralization, compared to other splenic macrophages. Collectively, our findings of ACE^+^ macrophages reflect the multitude of factors that shape macrophage phenotypes and their functions during persistent intracellular bacterial infection. They also illustrate how single-cell transcriptomics provide fuller pictures of cellular functional features that underlie the overall macrophage phenotypes and functional diversity.

## Materials and Methods

### Ethics Statement

Experiments involving animals were performed in accordance with NIH guidelines, the Animal Welfare Act, and US federal law. All animal experiments were approved by the Stanford University Administrative Panel on Laboratory Animal Care (APLAC) and overseen by the Institutional Animal Care and Use Committee (IACUC) under Protocol ID 12826. Animals were housed in a centralized research animal facility accredited by the Association of Assessment and Accreditation of Laboratory Animal Care (AAALAC) International.

### Mouse Strains and Husbandry

Females and males 129×1/SvJ mice were obtained from Jackson Laboratories or an in-house 129×1/SvJ colony. CD45.1^+^ 129×1/SvJ mice were generated from backcrossing CD45.1 ^+^ C57BL/6J mice (Jackson Laboratory) to 129×1/SvJ mice (Jackson Laboratory) successively for more than 10 generations. ACE 10/10 C57BL/6 mice were provided by Dr. Kenneth Bernstein. Male and female mice (7-16 weeks old) were housed under specific pathogen-free conditions in filter-top cages that were changed bi-monthly by veterinary or research personnel. Sterile water and food were provided *ad libitum*. Mice were given at least one week to acclimate to the Stanford Animal Biohazard Research Facility prior to experimentation.

### Bacterial Strains and Growth Conditions

*Salmonella enterica* serovar Typhimurium strain SL1344 was utilized in this study. SL1344 *ΔsteE*and SL1344 *Tomato* were generated as described previously(13, 28). For all mouse infections, *S*. Typhimurium strains were maintained aerobically on LB agar supplemented with 200 μg/mL streptomycin, ± 40 μg/mL kanamycin, and grown aerobically to stationary phase overnight at 37 °C with broth aeration. Bacterial cultures were spun down and washed with sterile phosphate-buffered saline (PBS) before suspension in PBS for infection.

### Mouse Infections and TNF neutralization

Mice were allocated to control and experimental groups randomly, sample sizes were chosen based on previous experience to obtain reproducible results and the investigators were not blinded. Mice were inoculated intraperitoneally (*i.p*.) with 1-2 × 10^3^ CFU *S*. Typhimurium SL1344 WT *or DsteE* in 200 mL PBS. For TNF neutralization, infected mice were injected *i.p*. with either 500 μg anti-TNF monoclonal Ab, clone MP6-XT22 (Biolegend), or isotype control Ab in sterile PBS in 400 μL total volume before scRNA-sequencing or analysis 4 days later. Mice were euthanized at the indicated time points post-inoculation by CO2 asphyxiation followed by cervical dislocation as the secondary method of euthanasia. Organs were collected, weighted, and either homogenized in PBS for CFU enumeration, used to make single cell suspension for flow cytometric analysis, or prepared for microscopy examinations.

### Bone-marrow chimera

CD45.2^+^ 129×1/SvJ mice at 6-8 weeks of age were lethally irradiated (600 Rad twice, 6 hours apart). Single-cell suspension from bone marrows of donor CD45.1^+^ 129×1/SvJ donor mice was prepared and injected intravenously into irradiated CD45.2^+^ mice via tail vein. Each recipient mouse received 1-2 × 10^^6^ of donor bone marrow cells. Recipient mice were maintained for 2 weeks on autoclaved food and water containing 2 mg/ml neomycin sulfate (VWR 89149–866) and 1000 U/ml polymyxin B (Millipore Sigma P4932-5MU). Bone marrow engraftment was assessed ~8 weeks after transplantation. Mice were infected with *S*. Typhimurium SL1344 9-10 weeks after transplantation and analyzed 1 month post-inoculation.

### Flow Cytometry

Spleens from mice were minced with surgical blades No. 22 and incubated in digestion buffer (HBSS + Ca^2+^ + Mg^2+^ + 50 μg/mL DNase (Roche) + 25 μg/mL Liberase TL (Sigma)) at 37 °C for 25 min, mixing at 200 rpm. EDTA was added at a final concentration of 5 mM to halt digestion. Single cell suspensions were passed through a 70-μm filter and washed with R5 buffer (RPMI containing 5% FBS and 10 mM Hepes). Red blood cells were lysed with ACK Lysis Buffer (Lonza) for 3 min at room temperature, washed, and resuspended in R5 buffer until they were stained for flow cytometry.

Single-cell suspensions were incubated in Fc Block (TruStain fcX anti-mouse CD16/32, Biolegend) for 15 min on ice and washed with PBS. Cells were stained on ice for 30 minutes in PBS with primary antibodies, followed by staining on ice for 30 minutes with a cocktail of Live/Dead Fixable Blue Viability Dye (Invitrogen) and fluorescent antibodies (list of antibodies used included). Cells were washed with FACS buffer (PBS containing, 2% FBS and 2 mM EDTA), followed by fixation for 15 min with Cytofix/Cytoperm solution (BD Biosciences). Cells were washed twice with Perm/Wash buffer (BD Biosciences) and stained for intracellular *Salmonella* and iNOS. After washing, cells were resuspended in a FACS buffer and analyzed on a LSRII cytometer (Becton Dickinson). Data were acquired with DIVA software (BD Biosciences) and analyzed using FlowJo software (TreeStar).

### Immunofluorescence Microscopy

Spleens were harvested, frozen in OCT compound (Fisher Scientific), and frozen sections 8 μm in thickness were placed on SuperFrost Plus cryosection slides (Fisher Scientific). Sections were fixed in ice-cold acetone at −20 °C for 10 minutes and then allowed to dry. A boundary was drawn around tissue sections using a pap pen (Fisher Scientific). Sections were washed with PBS and then blocked with a staining buffer (PBS with 3% bovine serum albumin and 5 % normal mouse serum) for 30 min at room temperature. After blocking, sections were stained with the primary antibodies in a staining buffer for 2 hours at room temperature. Sections were washed and then stained for 2 hours at room temperature with fluorescent conjugated secondary antibodies. Slides were washed in PBS and then mounted using ProLong Diamond (Life Technologies). Images were acquired on a Zeiss LSM 700 or 880 confocal microscope with the ZEN 2010 software (Zeiss) and processed using FIJI software.

### Cell preparation for 10X Genomics scRNA-sequencing

*S*Tm infected spleens were harvested and digested in buffer containing HBSS + Ca^2+^ + Mg^2+^ + 50 μg/mL DNase (Roche) + 25 μg/mL Liberase TL (Sigma) at 37 °C for 25 min, mixing at 200 rpm. EDTA was added to 5 mM final concentration to stop digestion reaction and cells were washed with RPMI containing 10% FCS. Following RBC lysis, splenocytes were washed twice RPMI containing 10% FCS. Splenocytes were stained with an antibody mixture for surface markers for 25 minutes on ice, washed twice with RPMI containing 10% FCS, then resuspended in the same buffer with 1:2000 DAPI. Splenocytes were then FACS-enriched on a BD FACSAria cell sorter. The viability of sorted cells were checked using Trypan blue staining and hemocytometer inspection under a light microscope. Samples had viability greater than 90%. Cells were resuspended to a concentration to 500-1200 cells/uL, partitioned, and captured for sequencing on a 10× Chromium Controller. Libraries were prepared by the Stanford Functional Genomics Facility (SFGF) using 10X Genomics 3’ GEX v3.1 kit and sequenced on the Illumina HiSeq4000 platform to a depth of ~40,000 - 50,000 reads/cell. Raw sequencing data were demultiplexed by SFGF to yield fastqs reads.

### Sequencing alignment and data preprocessing

Paired-end reads were mapped to *Mus musculus* genome reference GRCm38 using 10X cellranger (version 3.1.0) with parameters “--expect-cells=10000 --chemistry=auto”. Spliced and unspliced read counts were estimated by velocyto (v 0.17.17) in “run10x” mode. We performed downstream preprocessing and analyses on the UMI count matrices estimated by cellranger. To eliminate low-quality cells, we kept cells that had more than 500 detected genes and 1000 UMI read counts. We removed any stressed cells that had greater than 5% of total UMI counts that mapped to the mitochondrial genome. In total, we identified 22512 cells (WT +*ΔsteE S*Tm-infected mice) and 9892 cells (WT *S*Tm-infected mice treated with isotype control and anti-TNF antibodies) for downstream analysis. UMI counts were normalized for sequencing coverage such that each cell has a total number of counts equal to the median library size of all cells, yielding counts per median (CPM). The normalized CPM were added with a pseudocount of 1 and log2 transformed. Scanpy package (v 1.6.0) was used to perform data preprocessing and data transformation. RNA velocity was inferred using scVelo (v 0.2.2) with default parameters in “dynamical” mode.

### Cell clustering and cell type annotation

The SAM algorithm (version 0.8.5) was run with default parameters[36]. SAM outputs gene weights, principal component (PC) coordinates, a nearest-neighbor graph, and 2D UMAP projections used for visualization, on which the processed gene expression data can be also overlaid. We used the Leiden algorithm (version 0.8.4)(77) to determine the number of clusters in the entire immune population and inferred the immune cell type based on the approach proposed by panglaoDB(43). Briefly, a cell-type specific score was calculated for each cluster using a vector of cell-type specific genes and their associated weight:

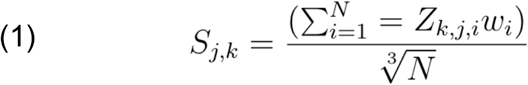

where S_j,k_ is the CTA score for cell-type j in cell cluster k and N is the total number of marker genes. Z is the Z-score standardized gene expression counts. For a given cell cluster, CTA scores are then ranked from highest to lowest and the top-ranking cell type is selected as the ‘winner’. We manually inspected the ‘winner’ cell type and determined that cluster 3 (Supplementary Figure 2B) was mis-annotated as “γΔT cells” since it did not express any T cell receptor γ, a definitive marker for γδ T cells. Because cluster 3 expresses mostly myeloid lineage genes, we annotated it as MNP 1.

### Differential gene expression analysis

Differential gene expression was computed with a negative binomial test (https://github.com/10XGenomics/cellranger/blob/master/lib/python/cellranger/analysis/diffexp.py). We compared the expression of each gene in each cluster against the rest of the populations. False discovery rate (FDR) was calculated using the Benjamini-Hochberg (BH) procedure. Genes were identified as differentially expressed genes (DEGs) based on FDR, log_2_ fold change in mean expression, and % detection in the query cluster.

### Myeloid cell, mononuclear phagocyte (MNP), and monocyte/macrophage sub-clustering

To ensure that we did not remove any of the MNPs (GMP, CM, or MΦ) for downstream analysis, we first sub-cluster the myeloid populations including Dendritic cells (DCs), Neutrophils, MNP 1, or MNP 2. We removed a few cells that appeared to be outliers (cell barcodes: ‘A1_CAGCGTGAGTCTAACC-1’, ‘A2_CACTGAAAGCGCCTCA-1’, ‘B2_GAGCTGCTCCGTGCGA-1’, ‘A3_ATACCGAGTTTCGACA-1’, ‘A3_CAATGACGTTGGCCTG-1’, ‘A3_GAAGGGTTCCGCACGA-1’, ‘A3_GAGTTTGTCAGCGTCG-1’, ‘A3_GTCTCACTCGGAATGG-1’, ‘B3_AACCCAAGTTGTCATG-1’, ‘B3_GATGGAGGTGCTCTCT-1’, ‘A4_CTACAGATCCACAGGC-1’, ‘A4_TAGGGTTCAAATGATG-1’, ‘B4_GTCACGGTCAGCCCAG-1’) as well as any cell that expressed Pou6f1 > 1 log_2_ CPM, a transcription factor typically detected in the mouse brain(78, 79). We removed genes that were detected > 1 UMI count in less than 20 cells and then preprocessed the data using SAMs preprocessing function with the parameters “norm=ftt”, “filter_genes=False”. We then executed the algorithm with the parameter “preprocessing=StandardScaler” to yield the myeloid sub-populations in Figure 1D. Leiden clustering was executed with parameter “resolution=0.9”. We manually annotated cell types based on the differentially expressed genes (DEGs) in each cluster.

### Mononuclear phagocyte (MNP) sub-clustering

To further analyze monocyte (CM) and macrophage (MΦ), we removed cells that were annotated as GMP in the MNP sub-populations (Leiden clusters 4, 8, 9, 10 in Figure 1D) and then preprocessed the data using SAMs preprocessing function with the parameter “filter_genes=False”. We executed the algorithm with default parameters to yield the monocyte/macrophage sub-populations (Figure 3A). Leiden clustering was executed with parameter “resolution=1.5”. We manually identified cell identities based on the expression level of myeloid marker genes and DEGs in each macrophage sub-population.

### Differential representation test on cell state

To test statistical association between infection conditions and cell state frequency, we stratified the dataset by experimental pairs (i.e., WT *S*Tm-infected mice 1 paired with *ΔsteE S*Tm-infected mice 1) and computed the chi-square distribution of counts for each condition and cell type. This ensures that we are contrasting cell state frequency in each condition while taking into account experimental variation due to sample processing or other technical factors. This test is also known as the Cochran-Mantel-Haenszel (CMH) test(80, 81). Briefly, we first prepared a series of 2 × 2 contingency tables (as many as there are experimental pairs) for each cell state:

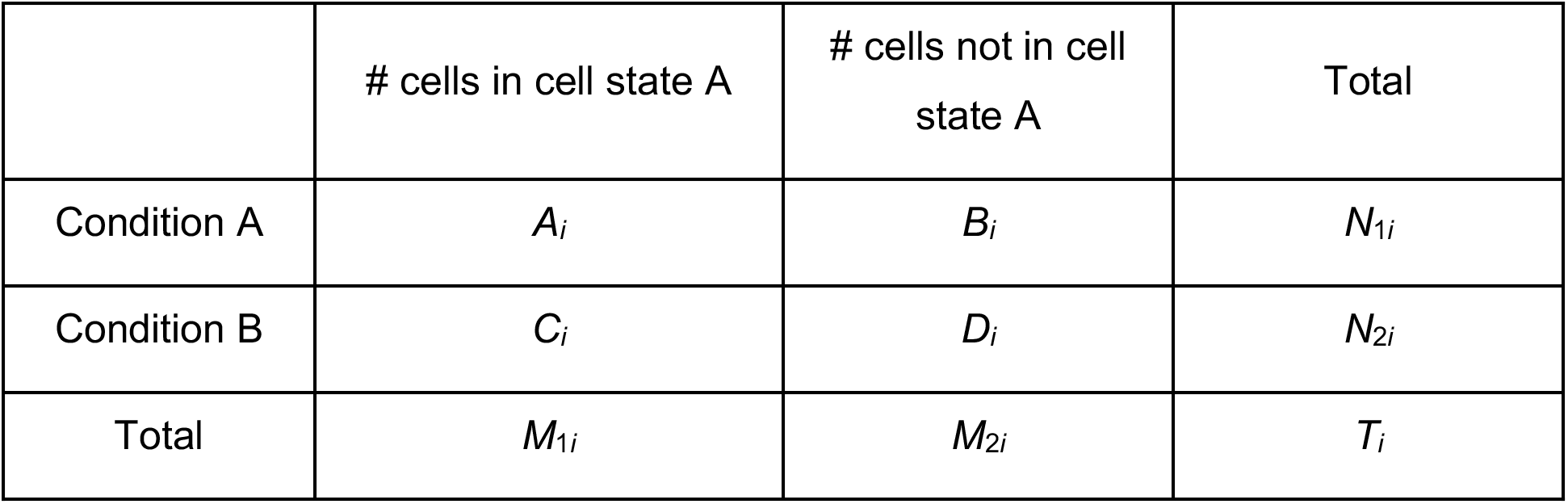

The common odds ratio is defined as:

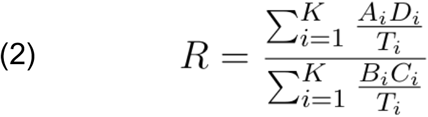

The test statistic is defined as:

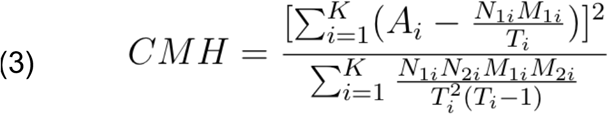

Where *i* is the index of the experimental strata (i.e., WT mice 1 vs *ΔsteE S*Tm-infected mice 1). Under the null hypothesis, there is no association between condition and cell state frequency for each stratum. The test statistic should then asymptotically follow a chi-square distribution. We accounted for multiple hypothesis testing by calculating the false discovery rate (FDR) with BH procedure. Cell type with greater than 2-fold difference in common odds ratio and less than 0.05 FDR between conditions were considered statistically significant.

### Gene sets over-representation analysis

To identify functional pathways that were enriched more than expected by chance, we performed gene set over-representation analysis (GSOA) on a list of DEG between cells of interest. Only DEGs that had FDR < 0.05, log_2_ fold change > 0.25, and > 25%detection were considered as input for GSOA. We measured the fraction of DEGs that belonged to each pathway under MSigDB Hallmark gene sets and computed the significance in overlap with hypergeometric test. The set size was set to the total number of mouse genes annotated in GRCm 38 (31,053 genes). FDR was calculated as before using BH procedure. Gene sets with FDR < 0.1 were considered statistically significant in enrichment.

### Ensemble expression score analysis

Score is calculated as per “score_genes” function in Scanpy with the provided gene list. Briefly, this is the average expression of the set of input genes subtracted by the average expression of a randomly sampled set of genes, controlled for expression range. A positive score indicates the average expression for the selected genes is above the group of randomly selected reference genes across the entire population.

### Cell cycle analysis and annotation

To predict the cell cycle phase of individual single cells, we curated a list of cell cycle related genes (S, and G2/M)(82). We then used the “score_genes_cell_cycle” function in Scanpy package to annotate the cell cycle phase of each cell.

## Acknowledgments

The authors thank Dr. Marc K. Jenkins, Department of Microbiology and Immunology, University of Minnesota Medical School, for providing the *S*. Typhimurium SL1344 *Tomato* strain. We thank members of the Denise Monack, Stephen Quake, and Manuel Amieva laboratories for valuable discussions. Research reported in this publication was supported by grant from the NIAID (TP), the Stanford Maternal and Child Health Research Institute (TP), the Stanford Pediatrics Department (TP), grant R01-AI116059 from the NIAID (DM), the Stanford Interdisciplinary Graduate Fellowship (YX), and the Chan-Zuckerberg Biohub (SQ).

## Funding

National Institute of Health grant K08-AI143796 (TP)

National Institute of Health grant R01-AI116059 (DM)

The Stanford Maternal and Child Health Research Institute (TP)

The Stanford Interdisciplinary Graduate Fellowship (YX)

The Chan-Zuckerberg Biohub (SQ)

## Author contributions

Conceptualization: THMP, DMM

Methodology: THMP, YX, SRQ, DMM

Investigation: THMP, YX, SMB, SRQ, DMM

Resource: KEB

Supervision: SRQ, DMM

Writing—original draft: THMP, YX, DMM

Writing—review & editing: THMP, YX, SMB, KEB, SRQ, DMM

### Competing interests

All authors declare no competing interests.

### Data and materials availability

Instructions to obtain processed data, preprocessing scripts, and analysis scripts are available on Github. Raw fastq files and processed data are deposited on SRA and GEO repository.

## Supplementary Materials

Available in supplementary file.

## Notes

### Competing Interest Statement

The authors have declared no competing interest.

