## Supplementary material for "Single-cell profiling identifies ACE^+^ granuloma macrophages as a non-permissive niche for intracellular bacteria during persistent *Salmonella* infection": SupplFigures

**This PDF file includes:**

Figs. S1 to S6

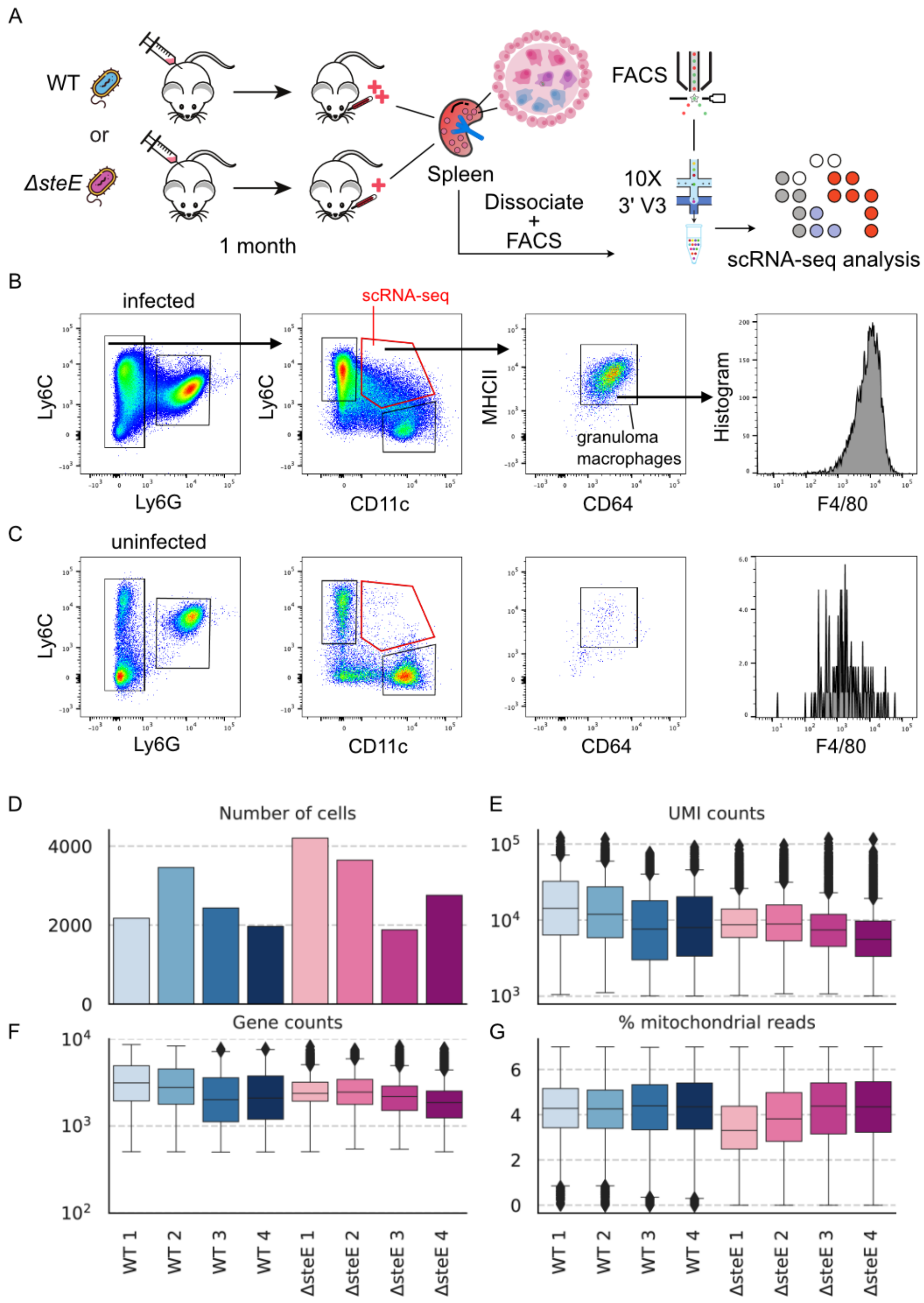


**Fig. S1.**

**Related to Figure 1 (A)** schematic outline of the droplet-based scRNA-seq of splenocytes from mice infected either with WT *S*Tm or *ΔsteE S*Tm. **(B-C)** FACS gating scheme for $CD11b^{+}CD11c^{+}Ly6C^{+}CD64^{+}MHC^{hi}$ granuloma macrophages, which also have high F4/80 expression. Samples were gated for size/scatter, singlet, living CD11b$^+$ population then further gated as displayed. Plots shown are results from WT-*S*Tm infected. For permissive FACS enrichment of CD11b$^+$CD11c$^+$Ly6C$^+$ granuloma macrophages, sorting gates were set tightly for size/scatter, singlet, and living cells but more loosely for CD11b, Ly6G, Ly6C, and CD11c markers. This enrichment strategy enables significant enrichment of macrophages of granuloma phenotypes and simultaneous capture the full spectrum of immune cells in infected spleens. Red arrow indicates the CD11b$^+$CD11c$^+$Ly6C$^+$ mononuclear phagocyte (MNP) population targeted for enrichment for scRNA-seq library preparation. **(D)** Number of analyzed cells that passed the quality controls from individual mice. Cells that had < 1000 unique molecular identifier (UMI) counts, < 500 detected genes, and > 5% reads with mitochondrial origin (indicate stressed or dying cells) were filtered. A total of 22,512 cells from WT and *ΔsteE*-*S*Tm infected mice passed quality controls. **(E)** Distribution of UMI counts in analyzed cells. **(F)** Distribution of the number of detected genes in analyzed cells. **(G)** Distribution of the percent reads that mapped to mitochondrial genes in analyzed cells.

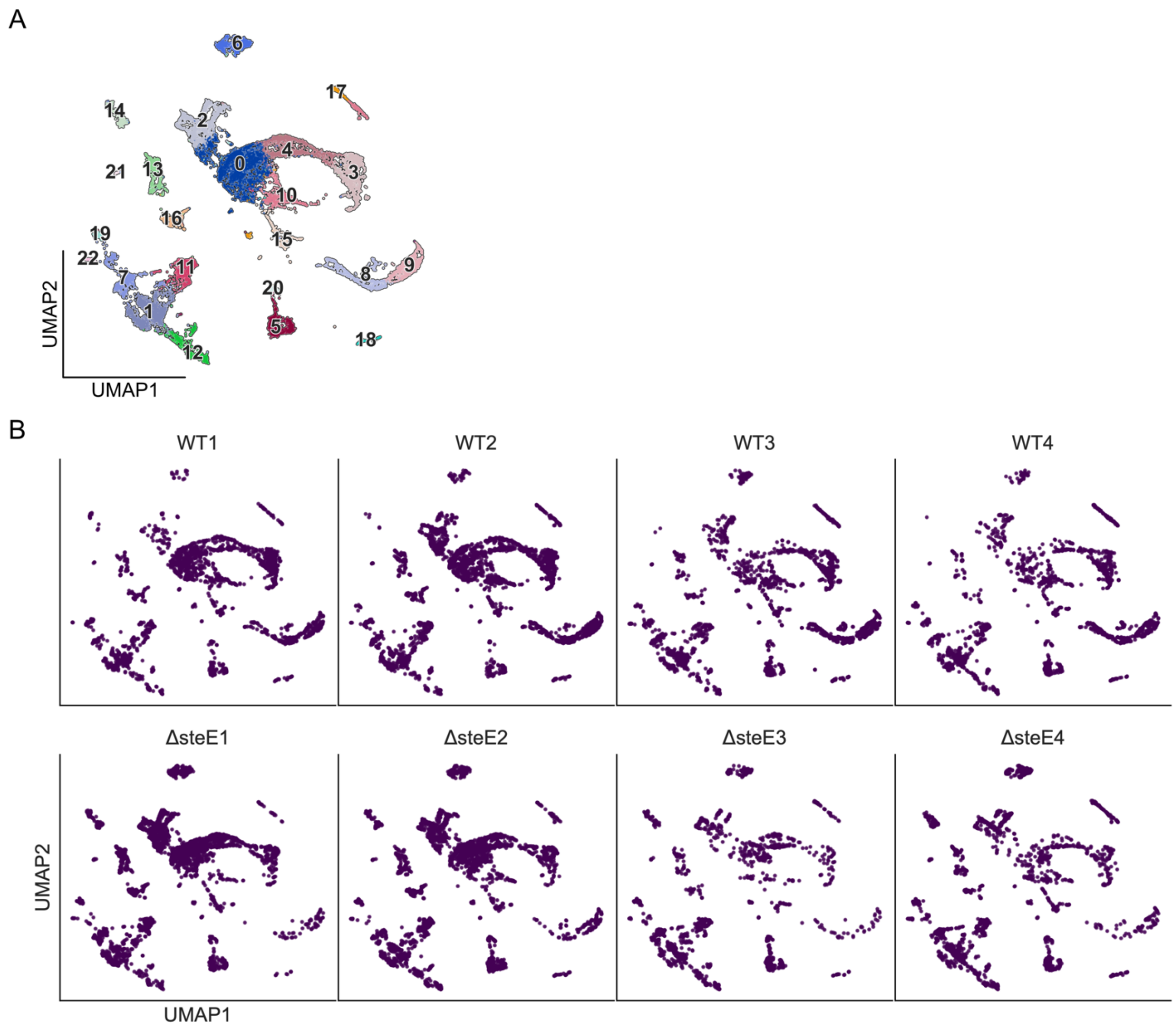


**Fig. S2.**

**Related to Figure 1. (A)** UMAP projection colored by Leiden clusters. **(B)** UMAP projections of cells from either WT *S*Tm or *ΔsteE S*Tm-infected mice. Cells are well mixed in each part of the manifold, indicating a lack of technical batch effect.

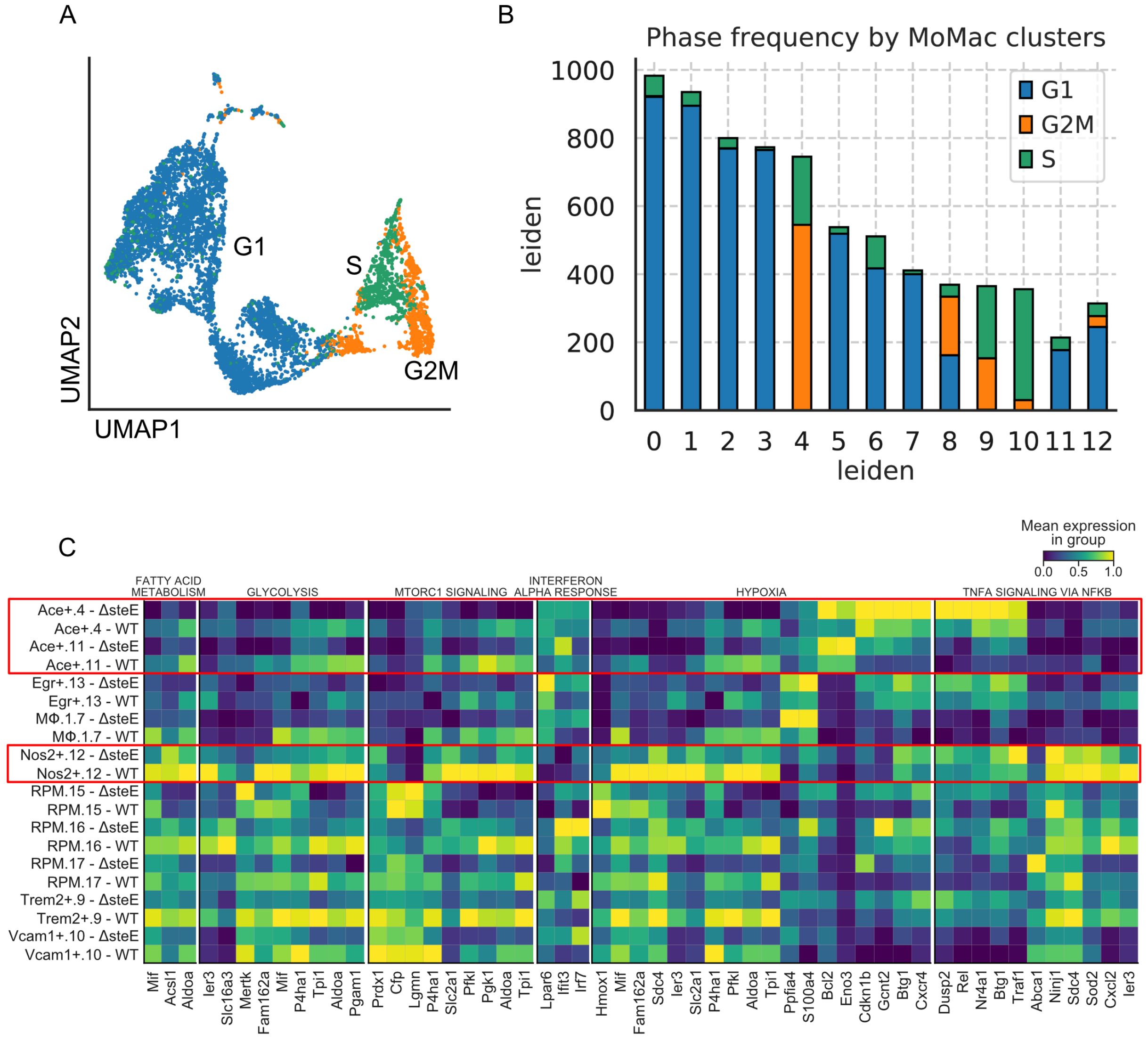


**Fig. S3.**

**Related to Figure 3. (A)** UMAP projection of myeloid populations colored by predicted cell cycle states. **(B)** Cell cycle phase frequency for each MNP cluster. **(C)** Heatmap of standardized expression of genes in selected pathways in Figure 3H.

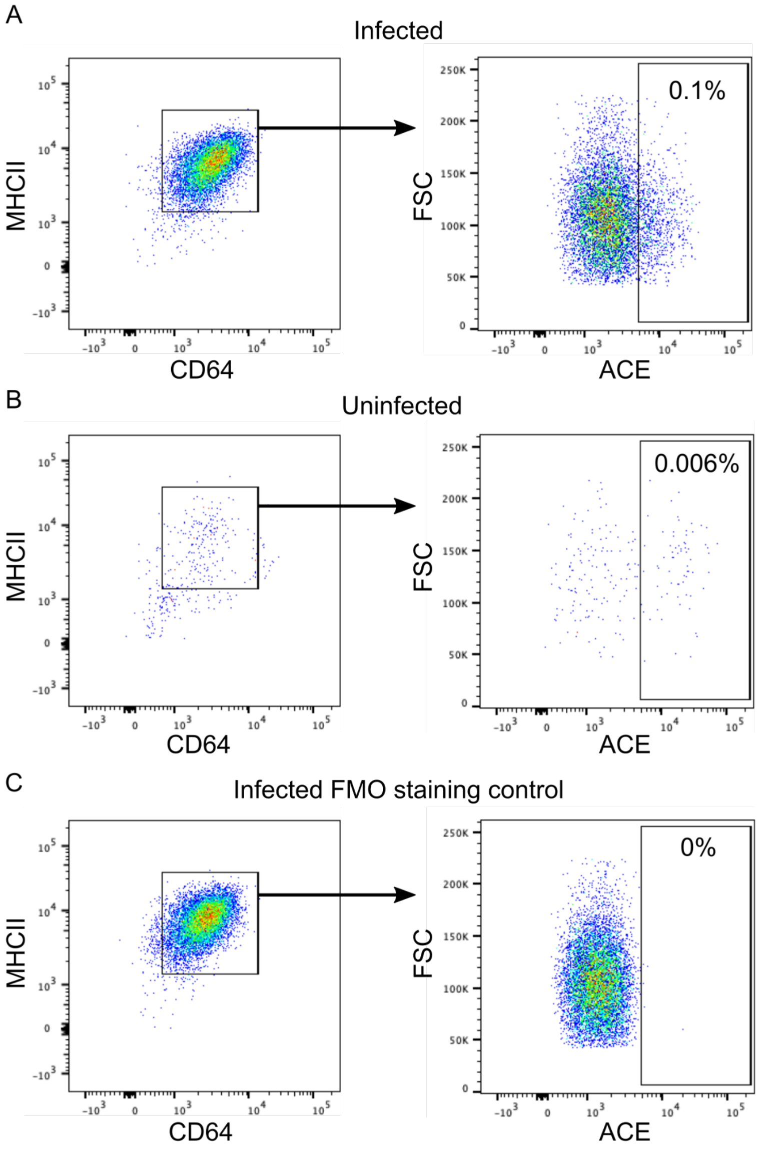


**Fig. S4.**

**Related to Figure 4.** Mice were infected intraperitoneally with 2 x $10^3$ CFU WT-*S*Tm and analyzed at 1 month post-inoculation. Flow cytometry gating for $ACE^+$ granuloma macrophages. Splenocytes were gated for singlet, living $CD11b^+CD11c^+Ly6C^+$ population, then gated as displayed. Percentages indicate frequencies of gated cells among total splenocytes. Plots shown are results from infected mice **(A)**, uninfected mice **(B)**, or infected mice but without ACE antibody stain as a control **(C)**.

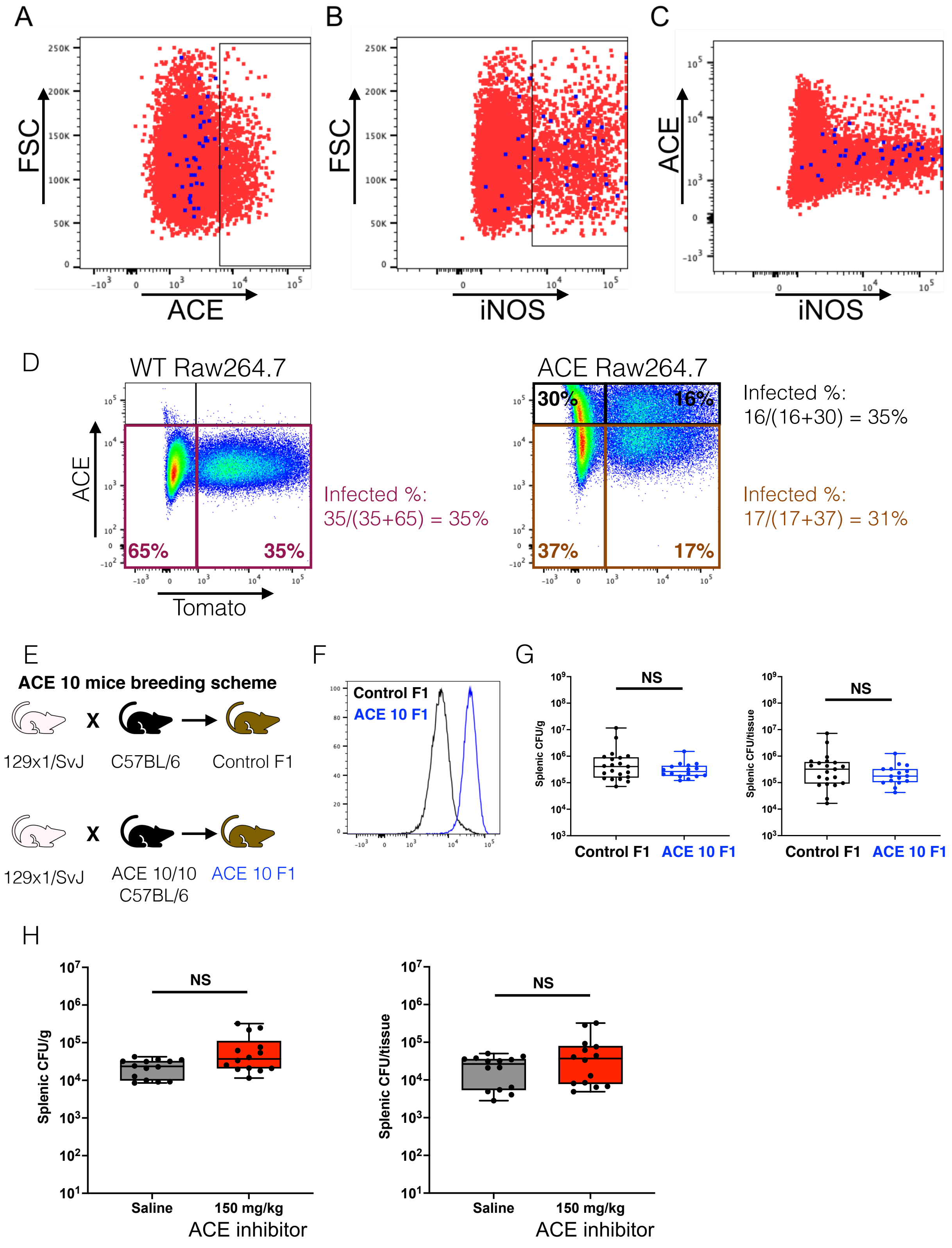
A
FSC
ACE
B
FSC
iNOS
C
ACE
iNOS
D
WT Raw264.7
ACE Raw264.7
ACE
Tomato
65%
35%
Infected %:
35/(35+65) = 35%
30%
16%
37%
17%
Infected %:
16/(16+30) = 35%
Infected %:
17/(17+37) = 31%
E
ACE 10 mice breeding scheme
129x1/SvJ
C57BL/6
Control F1
129x1/SvJ
ACE 10/10
C57BL/6
ACE 10 F1
F
Control F1
ACE 10 F1
G
NS
Splenic CFU/g
Control F1
ACE 10 F1
NS
Splenic CFU/tissue
Control F1
ACE 10 F1
H
NS
Splenic CFU/g
Saline
150 mg/kg
ACE inhibitor
NS
Splenic CFU/tissue
Saline
150 mg/kg
ACE inhibitor

**Fig. S5.**

**Related to Figure 5. (A-C)** Mice were infected *intraperitoneally* (*i.p.*) with 2 x $10^3$ CFU WT-*S*Tm and analyzed at 1 month post-inoculation. Flow cytometry showing *S*Tm-containing cells among $iNOS^+$ and $ACE^+$ granuloma macrophages. $CD11b^+CD11c^+Ly6C^+CD64^+MHC^{hi}$ granuloma macrophages were gated as in Figure S1B, then plotted for FSC vs. ACE (A), FSC vs. iNOS (B), or ACE vs. iNOS (C) as shown. Red events: all granuloma macrophages, blue events: granuloma macrophages harboring intracellular *S*Tm. **(D)** *Ace* was expressed in RAW264.7 macrophages using lentiviral transduction. WT and ACE-overexpressing RAW264.7 macrophages were infected with *S*Tm and analyzed by flow cytometry 20 hours later. The percent frequencies of $STm^+$ cells were similar between $ACE^-$ and $ACE^+$ cells. **(E)** Breeding scheme to generate [ACE 10/10 x 129x1/SvJ] F1 mice and C57BL/6 x 129x1/SvJ F1 control mice for infection. **(F-G)** Mice were infected with 1-2 x $10^3$ CFU WT-*S*Tm *i.p.* and analyzed at two weeks post-inoculation. **(F)** Flow cytometry analysis shows splenic $CD11b^+CD11c^+Ly6C^+CD64^+MHC^{hi}$ granuloma macrophages from infected ACE 10 heterozygous F1 mice expressing significantly higher ACE levels compared to granuloma macrophages from control F1 mice. **(G)** CFU measurements shows similar *S*Tm bacterial levels in the spleens of ACE 10 heterozygous F1 mice and control mice. **(H)** Inhibition of ACE enzymatic activity *in vivo* had no significant impact on *S*Tm levels in infected spleens. Mice were infected for 1 month then treated with either saline control or 150 mg/kg of the ACE inhibitor captopril *i.p.* daily for 7 days before analysis. **A**, **H**. Dot: individual mice.

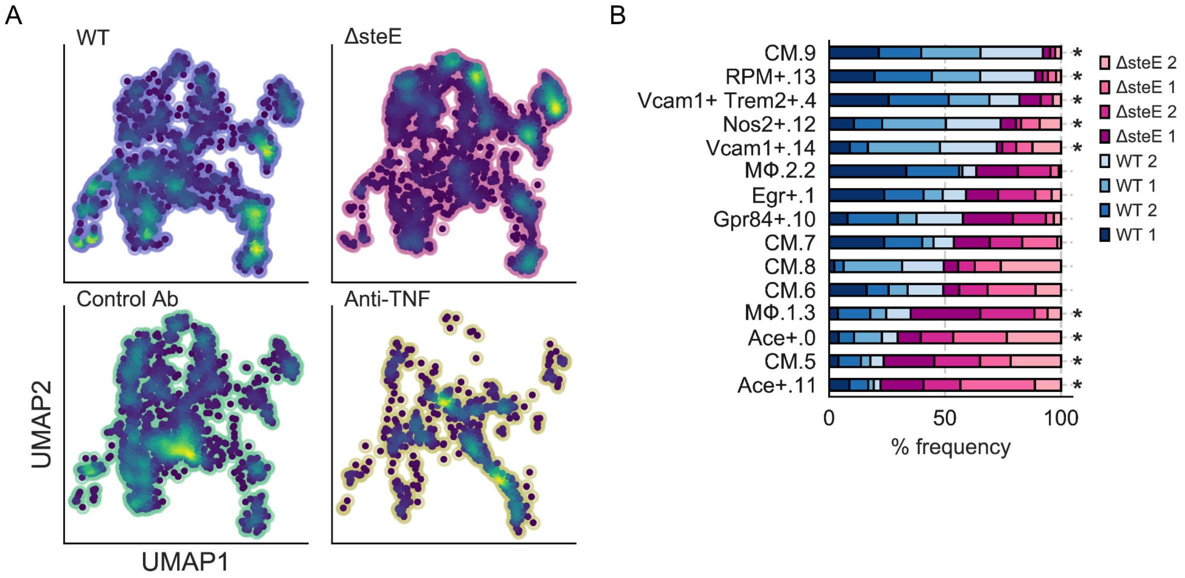


**Fig. S6.**

**Related to Figure 6. (A)** UMAP projections of mononuclear phagocytes, excluding dendritic cells, isolated from WT *S*Tm, *ΔsteE S*Tm-infected mice, or WT *S*Tm-infected mice treated with control antibody or anti-TNF. Color reflects cell density in each part of the manifold. **(B)** Differential representation of cells. Clusters were defined based on co-clustering with cells from WT *S*Tm, *ΔsteE S*Tm-infected mice, or WT *S*Tm-infected mice treated with control antibody or anti-TNF. Asterisk next to the bar indicates a greater than 2-fold difference in representation ratio and statistical significance between WT *S*Tm and *ΔsteE S*Tm infection based on a differential representation test (FDR < 0.05; see Methods).
